# Optimized dosing of spatially heterogeneous radiopharmaceutical therapy reprograms poorly immunogenic tumors to enhance response to immune checkpoint blockade

**DOI:** 10.64898/2026.10.27.762270

**Authors:** Maya E. Takashima, Zachary Danaceau, Malick Bio Idrissou, Ohyun Kwon, Tracy J. Berg, Paul A. Clark, Yujuan Wang, Wonjong Jin, Adedamola O Adeniyi, Campbell Donnelly Haasch, Michael Antonelli, Jamey Weichert, Bryan Bednarz, Reinier T. Hernandez, Irene Ong, Zachary S. Morris

**Affiliations:** Department of Radiation Medicine, University of Wisconsin School of Medicine and Public Health, University of Wisconsin-Madison, Madison, WI, USA 53705; Department of Biostatistics and Medical Informatics, University of Wisconsin School of Medicine and Public Health, University of Wisconsin-Madison, Madison, WI, USA 53705; Department of Medical Physics, University of Wisconsin School of Medicine and Public Health, University of Wisconsin-Madison, Madison, WI, USA 53705; Department of Radiology, University of Wisconsin School of Medicine and Public Health, University of Wisconsin-Madison, Madison, WI, USA 53705

## Abstract

Immune checkpoint inhibitors (ICIs) are ineffective in patients with poorly immunogenic tumors due to exclusion or exhaustion of effector T cells in the tumor microenvironment (TME) and/or inadequate priming of tumor-specific T cells. Preclinical and clinical studies suggest that radiopharmaceutical therapy (RPT) may overcome such challenges, but an optimal approach to RPT dosing has not been clarified in combination with ICIs. Conventional RPT dosimetry emphasizes mean absorbed radiation doses in tumors and normal tissues, neglecting the spatial distribution of this dose in the TME. We hypothesize that RPT dose heterogeneity in tumors may be leveraged to initiate spatially restricted immuno-radiobiological effects in TME that potentiate antitumor immunity when injecting an activity that delivers high dose (V_10%_>20 Gy) while also preserving low dose regions (V_90%_<2.5 Gy) in the TME and not inducing systemic lymphopenia. In studies using lutetium-177 RPT in combination with ICIs, administered activities that met these dosimetric optimization metrics occurred at different mean tumor doses for different RPT agents and different models, yet each achieved therapeutic efficacy in poorly immunogenic murine tumors. Spatial transcriptomics at day 7 after RPT revealed heterogeneous distribution of tumor cell markers with distinct dose-response profiles and localization of dendritic cell (DC) migratory markers in regions of moderate/high dose response marker expression. Combination therapy sustained conventional type 1 DC (cDC1) activation in tumor draining lymph nodes and promoted systemic expansion of antigen-specific CD8 T cells, while serial biopsies revealed effector T cell recruitment and inflammatory cytokine expression in the TME. Optimized intratumoral dose heterogeneity may enable and predict response to ICIs when combined with RPT in tumors not responding adequately to ICIs alone.

## INTRODUCTION

Immune checkpoint inhibitors (ICIs), such as anti-PD-L1 and anti-CTLA-4, function by disrupting inhibitory immune signaling pathways that regulate T-cell priming and effector activity^1^. Patients responding to ICIs can experience complete and durable control of metastatic disease, however for most patients clinical efficacy is limited by primary or acquired resistance^2^. Because effective checkpoint blockade depends on the generation and recruitment of primed tumor-reactor T cells, blocking inhibitory pathways alone may remain ineffective in tumor microenvironments (TME) lacking effective antigen presentation and functional tumor-infiltrating T cells^3,4^. Overcoming ICI resistance requires strategies that actively induce antigen presentation and convert non-responsive TMEs into receptive immune landscapes.

Radiopharmaceutical therapy (RPT) offers a compelling strategy to modulate the TME by delivering systemic radiation to multiple sites of disease^5–7^. Beyond direct tumor cell killing, radiation serves as a versatile dose-dependent immune modulator^8–11^. High-dose radiation can promote immunogenic cell death, releasing damage-associated molecular patterns and tumor antigens necessary for antigen-presenting cell recruitment^10,12–14^. Moderate-dose radiation can stimulate type I interferon signaling, while low-dose radiation may promote endothelial adhesion and immune cell infiltration^10,12,15–17^. Despite these dose-dependent effects, the selection of RPT doses for combination immunotherapy has largely focused on administered activity or mean tumor absorbed dose rather than the spatial distribution of absorbed dose within the TME^18,19^. Rooted in radiobiological paradigms aimed at maximizing tumor cell killing by tumor cell DNA damage, escalating RPT dose may reduce intratumoral low-dose regions and compromise local or systemic immune populations, potentially blunting the anti-tumor immune response that is also a component of response to RPT and that is critical for durable response to ICIs^10,12^.

Building on our previous finding that certain key metrics of RPT dose heterogeneity can predict therapeutic response in a monotherapy setting through immune activation^10^, here we test whether these same metrics of spatially-resolved dosimetry can predict immuno-radiobiological effects and identification of an effective RPT dosing approach when combined with ICIs. While non-uniform dosing has traditionally been considered a therapeutic limitation for RPT, our prior data suggests that this dose heterogeneity may present an opportunity for mitigating ICI resistance by enabling preservation of low-dose regions in the TME that facilitate immune cell survival and function while simultaneously promoting immune cell activation and tumor cell immune susceptibility in moderate and high dose regions. We hypothesize that, by selecting an administered activity of lutetium-177 RPT that delivers the highest tumor dose possible while not inducing systemic lymphopenia and while maintaining a maximum dose of <2.5 Gy in >10% of the tumor, we will activate the conventional type I dendritic cell (cDC1) – CD8 T cell priming axis and enable anti-tumor immunity in tumors not responding to dual ICI therapy.

Here, we test this hypothesis and evaluate the therapeutic mechanisms of integrating a spatially heterogeneous dose of lutetium-177 into dual checkpoint blockade treatment (anti-PD-L1 and anti-CTLA-4). We first investigate the utility of image-based dosimetry for predicting dose optimization of ^177^Lu-PNT6555, targeting FAP^20^, in mice bearing mixed B16-mFAP^+^/mFAP^−^ melanoma. We extend this dosing strategy to the immunologically cold MOC2 head and neck squamous cell carcinoma and KP sarcoma models using ^177^Lu-NM600^21^. Flow cytometry on disaggregated tumor and tumor-draining lymph node (TDLN), TME secretome profiling, and spatial transcriptomics are employed to evaluate immune infiltration, functional responses, and dose-dependent gene signatures. Our preclinical findings support our hypothesis, demonstrating that dosing of RPT to achieve maximal tumor dose without inducing lymphopenia or eliminating low dose regions in the TME achieves DC activation and migration, enhances antigen-specific CD8 T cell response, and promotes an effector-recruiting secretome in the TME. Furthermore, we demonstrate that continued post-RPT checkpoint blockade is important for preserving this modified immune state, establishing a rationale for integrating spatially informed RPT dose selection with optimized immunotherapy sequencing in ICI-refractory cancers.

## RESULTS

### Heterogeneous dose of ^177^Lu RPT in combination with dual immune checkpoint inhibition leads to longer overall survival

To evaluate whether an immunologically optimized heterogeneous dose of ^177^Lu RPT enhances sensitivity to ICI, we used our previously published dosimetric modeling^10^ to test whether metrics of RPT dose heterogeneity that we identified as predictive of response to RPT monotherapy also predicted therapeutic response in combination with ICIs. For this, we tested mean tumor doses of 2.5, 5, 10, or 20 Gy from ^177^Lu-PNT6555 in orthotopic murine melanoma skin tumors composed of B16-mFAP^+^ and B16-mFAP^−^ cells. Cumulative dose-volume histograms (DVH) derived from dosimetric modeling and supported by *ex vivo* ionizing-radiation quantum imaging detector (iQID) indicate that a mean absorbed tumor dose of 2.5 Gy from ^177^Lu-PNT6555 in this model does not achieve localized high dose (>20 Gy) regions in the TME while mean tumor doses of 10 and 20 Gy do not preserve regions of tumor getting below 2 Gy (Figure 1A and 1B). 5 Gy ^177^Lu-PNT6555 preserves low dose regions while also achieving regions of high dose (Figure 1B). We tested whether this dose would be optimal in combination with dual ICI (anti-CTLA4 and anti-PD-L1). C57BL/6 mice bearing mixed B16-mFAP^+^/mFAP^−^ tumors were randomized to no treatment, dual ICI alone, 2.5 Gy ^177^Lu-PNT6555 + dual ICI, 5 Gy ^177^Lu-PNT6555 + dual ICI, 10 Gy ^177^Lu-PNT6555 + dual ICI, or 20 Gy ^177^Lu-PNT6555 + dual ICI. Mice receiving dual ICI were intraperitoneally injected with 200 μg of anti-CTLA4 and anti-PD-L1 on day −3, 0, 3 with RPT on day 1. An injected activity of ^177^Lu-PNT6555 that delivered a mean tumor dose of 5 Gy in combination with dual ICI led to a significant improvement in overall survival compared to all other groups (p<0.05; Figure 1C), consistent with our hypothesis that, when combined with ICI therapy, the optimal injected activity of an RPT may be that which maximizes tumor dose while not inducing systemic lymphopenia and while maintaining a maximum dose of <2 Gy in >10% of the tumor.

**Figure 1.**
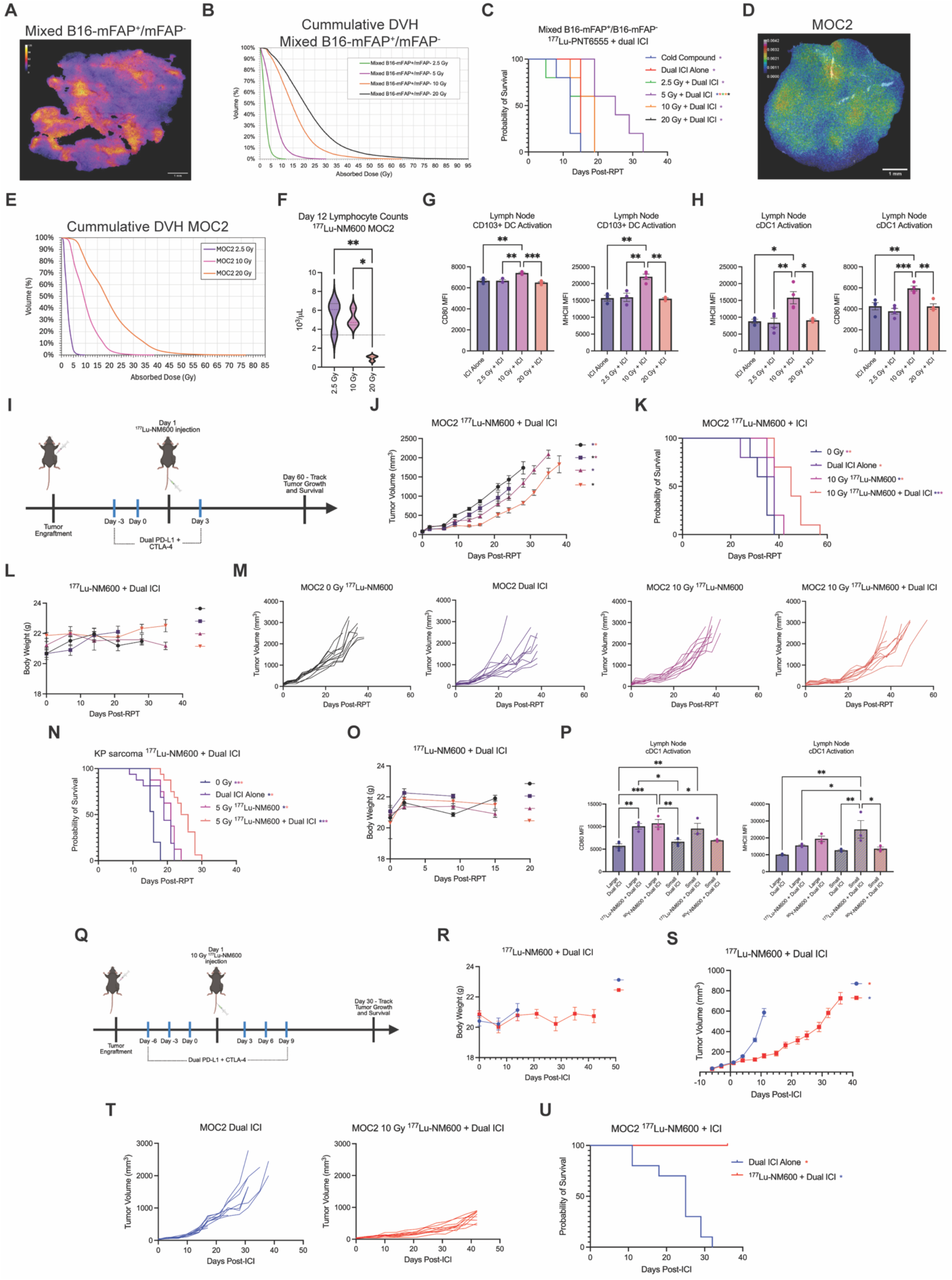
Dose heterogeneity from ^177^Lu-NM600 improves response to dual immune checkpoint inhibition. A) Activity maps from ionizing-radiation quantum imaging detector for mixed B16-mFAP^+^/mFAP^−^ at 24h post injection of 666 μCi ^177^Lu-PNT6555. B) Cumulative DVH curves for mixed B16-mFAP^+^/mFAP^−^ tumors receiving 2.5, 5, 10, or 20 Gy ^177^Lu-PNT6555. C) Mixed B16-mFAP^+^/mFAP^−^ tumor-bearing mice were randomized to no treatment, dual ICI (anti-PD-L1 + anti-CTLA-4), 2.5 Gy ^177^Lu-PNT6555 + dual ICI, 5 Gy ^177^Lu-PNT6555 + dual ICI, 10 Gy ^177^Lu-PNT6555 + dual ICI, or 20 Gy ^177^Lu-PNT6555 + dual ICI. D) Activity maps from ionizing quantum imaging detector for MOC2 at 24h post injection of 256 μCi ^177^Lu-NM600. E) Cumulative DVH curves for MOC2 tumors receiving 2.5, 10, or 20 Gy ^177^Lu-NM600. F) Lymphocyte count at day 12 following injection of ^177^Lu-NM600. G-H) Analysis of CD80 and MHC class II MFI in tumor draining lymph node (TDLN) type 1 conventional dendritic cells, cDC1 (CD45+CD11b+CD11c+MHCII+XCR1+) and CD103+ DCs (CD45+CD11b+CD11c+CD103+) at day 12 in MOC2 tumor-bearing mice treated with dual ICI alone, 2.5 Gy ^177^Lu-NM600 + dual ICI, 10 Gy ^177^Lu-NM600 + dual ICI, or 20 Gy ^177^Lu-NM600 + dual ICI. I) Schematic for treatment timing. J-M) MOC2 tumor-bearing mice were randomized to no treatment, dual ICI (anti-PD-L1 + anti-CTLA-4), 10 Gy ^177^Lu-NM600, or 10 Gy ^177^Lu-NM600 + dual ICI. Dual ICI was administered on day −3, 0, 3 with ^177^Lu-NM600 administered on day 1. J-M) Effect of combination 10 Gy ^177^Lu-NM600 and dual ICI. N-O) KP sarcoma tumor bearing mice were randomized to no treatment, dual ICI (anti-PD-L1 + anti-CTLA-4), 5 Gy ^177^Lu-NM600, or 5 Gy ^177^Lu-NM600 + dual ICI. Dual ICI was administered on day −3, 0, 3 with ^177^Lu-NM600 administered on day 1. N-O) Effect of combination 5 Gy ^177^Lu-NM600 and dual ICI. P) Analysis of CD80 and MHC class II MFI in tumor draining lymph node (TDLN) type 1 conventional dendritic cells, cDC1 (CD45+CD11b+CD11c+MHCII+XCR1+) at day 7 in large and small KP sarcoma-tumor bearing mice treated with dual ICI alone, 5 Gy ^177^Lu-NM600 + dual ICI, or 5 Gy ^90^Y-NM600 + dual ICI. Q) Schematic for treatment timing. R-U) MOC2 tumor-bearing mice were treated with dual ICI (anti-PD-L1 + anti-CTLA-4) or 10 Gy ^177^Lu-NM600 + dual ICI. Dual ICI was administered day −6, −3, 0, 3, 6, and 9 with ^177^Lu-NM600 administered on day 1. Blood was collected via submandibular vein at day 0 and 12. R-U) Effect of prolonged ICI treatment in combination with 10 Gy ^177^Lu-NM600. A-U) Injected activities were as followed: mixed B16-mFAP^+^/mFAP^−^ 2.5 Gy ^177^Lu-PNT6555=167 μCi; mixed B16-mFAP^+^/mFAP^−^ 5 Gy ^177^Lu-PNT6555=340 μCi; mixed B16-mFAP^+^/mFAP^−^ 10 Gy ^177^Lu-PNT6555=667 μCi; mixed B16-mFAP^+^/mFAP^−^ 20 Gy ^177^Lu-PNT6555=1,333 μCi; MOC2 2.5 Gy ^177^Lu-NM600=64 μCi; MOC2 10 Gy ^177^Lu-NM600=256 μCi; MOC2 10 Gy ^177^Lu-NM600=513 μCi; KP sarcoma 5 Gy ^177^Lu-NM600=251 μCi; KP sarcoma 5 Gy ^90^Y-NM600= 113 μCi C) N=5 all groups. F-H) N=3 all groups. I-L) N=10 all groups. M) N=3-5 all groups. N-O) N=15 all groups. P) N=3 all groups. R-U) N=9-10 all groups. Linear mixed model was used to compare tumor growth and log-rank test was used to compare survival. *p < 0.05 by linear mixed model (J and S), by Kaplan-Meier method (C, K, U) the color of the asterisk represents the group from which the group differs, *p < 0.05, **p < 0.01, ***p < 0.001, ****p < 0.0001 by one-way ANOVA (G, H, P).

Applying this predictive approach in a separate poorly immunogenic model and with a different RPT agent, we next evaluated ^177^Lu-NM600 in mice bearing MOC2 murine head and neck squamous cell carcinoma. Our dosimetric model and complete blood counts identified an injected activity of ^177^Lu-NM600 that delivered 10 Gy mean tumor dose as the highest injected activity that would not cause lymphopenia and that preserved low-dose regions in the MOC2 TME (Figure 1D-F). We previously identified dendritic cell (DC) preservation and functional activation as a critical immunologic effect of heterogeneous radiation dosing that is dependent upon maintaining regions in the TME that only receive low dose^10,12^. To evaluate whether a 10 Gy mean tumor dose from ^177^Lu-NM600 in MOC2 tumors was optimal to activate DCs in combination with dual ICI we compared dual ICI alone, 2.5 Gy + ICI, 10 Gy + ICI, and 20 Gy + ICI in mice bearing MOC2. 10 Gy ^177^Lu-NM600 in combination with dual ICI led to a significant increase in CD80 and MHCII MFI on conventional type I dendritic cells (cDC1s) at day 12 after RPT administration, as compared to ICI alone, 2.5 Gy + ICI, and 20 Gy + ICI (CD80: p<0.01 and MHCII: p<0.05; Figure 1G and 1H). To test the therapeutic effect of combining 10 Gy ^177^Lu-NM600 and dual ICI, we engrafted C57BL/6 with MOC2 and randomized mice to no treatment, 10 Gy ^177^Lu-NM600 alone (via tail vein), dual ICI alone, or 10 Gy ^177^Lu-NM600 + dual ICI (Figure 1I-M). Compared to no treatment, dual ICI alone significantly reduced tumor growth (p=0.002, Figure 1J) but did not improve overall survival. An immune-optimized dose of ^177^Lu-NM600 alone significantly improved overall survival (p=0.004, Figure 1K). Mice treated with combination of an immune-optimized dose of ^177^Lu-NM600 + dual ICI had a significant reduction in tumor growth compared to no treatment (p=0.011, Figure 1J) and a significant extension in overall survival compared to dual ICI alone or RPT alone (p=0.001; Figure 1K).

To further corroborate the effective application of this approach to optimizing RPT dosing in combination with ICIs, we used the poorly immunogenic KP soft tissue sarcoma model engrafted orthotopically in the right thigh musculature. Using prior dosimetric studies^10^, we identify a 251 µCi injected activity of ^177^Lu-NM600 delivering 5 Gy mean tumor dose as the highest injected activity that preserved low-dose regions in the KP TME. We tested the anti-tumor efficacy of combining this 5 Gy dose of ^177^Lu-NM600 with dual ICI in SvJ mice bearing KP sarcoma. Mice were randomized to no treatment, ^177^Lu-NM600 alone, dual ICI alone, or ^177^Lu-NM600 + dual ICI (Figure 1N-O). Because of the deep implantation of these tumors, response and metrics for euthanasia were based on morbidity metrics as surrogates for cancer progression (Table 2). Mice were euthanized when a Grade 4 toxicity was observed. Compared to no treatment, both ICI alone and ^177^Lu-NM600 alone had a significant improvement in overall survival (p<0.001, Figure 1N). Notably the combination ^177^Lu-NM600 + ICI led to significant improvement in overall survival compared to ^177^Lu-NM600 alone (p<0.001) and ICI alone (p<0.001; Figure 1N). These data support our hypothesis and indicate that an optimized dose of RPT can enhance response to dual ICI therapy and improve survival in mice bearing poorly immunogenic tumors that are resistant to ICIs alone.

Because the path length of radioisotope emissions and the physical size of a tumor fundamentally alter the distribution of dose in the TME following treatment with an RPT agent, we sought to compare the activation of DCs in the tumors of mice bearing orthotopically implanted KP sarcoma following treatment with ^177^Lu-NM600 or ^90^Y-NM600. The median range of β-particle emission from ^177^Lu is 0.67 mm, whereas that from ^90^Y is 2.5 mm. We therefore hypothesized that ^177^Lu-NM600 would better preserve low dose regions in the TME for a given mean tumor dose in small tumors, whereas in large tumors this effect might wane as the physical size of stromal regions in tumors begins to exceed mean β-particle range. Evaluating cDC1 activation in KP sarcoma-bearing mice, we observed that while both ^177^Lu-NM600 and the longer-range beta emitter ^90^Y-NM600 significantly increased cDC1 CD80 and MHCII MFI in large tumors (diameter ∼ 16mm), whereas only ^177^Lu-NM600 maintained robust cDC1 activation in small tumors (diameter ∼ 5mm; p<0.05; Figure 1P). In contrast, ^90^Y-NM600 failed to activate cDC1s in small tumors, suggesting that shorter range RPT emissions may be required to achieve advantageous effects of dose heterogeneity on anti-tumor immune response in small tumors.

Given the modest therapeutic response seen with three doses of ICI concurrent with RPT, we examined whether adding RPT to a regimen that included ICI treatment both prior to and after RPT, would improve survival and anti-tumor immune response. We administered three pre-treatment dual ICI doses (Days 0, 3, 6) prior to ^177^Lu-NM600 on Day 7, followed by three post-treatment dual ICI doses (Day 9, 12, 15) in MOC2-bearing mice (Figure 1Q). Body weight was not significantly affected by the treatment regimen (Figure 1R). Mice bearing established MOC2 tumors were randomized to either dual ICI alone or 10 Gy ^177^Lu-NM600 + dual ICI, and tumor growth and survival was tracked for 30 days following ^177^Lu-NM600 injection (Figure 1S-U). While tumors in mice receiving dual ICI alone continued to progress, combination therapy significantly delayed tumor growth (p=0.004; Figure 1S-T). Consequently, combination treatment significantly improved overall survival (p<0.0001; Figure 1U), maintaining 100% survival at 30 days post-RPT compared to a median survival of 25 days in mice receiving dual ICI alone.

### RPT modifies the TME secretome, a predictive biomarker of response to ICIs

Functional changes in chemokines and cytokine production after irradiation may contribute to immune cell recruitment and activation in the TME^22,23^. Given persistence of DC and CD8 T cell responses after RPT and continued ICI treatment, we evaluated secretome changes in the TME by culturing live *ex vivo* core biopsies on the Elephas platform (Figure 2A)^24^. The Elephas platform utilizes a sequential *ex vivo* treatment of live tumor fragments to compare ICI response against baseline isotype control within the same tissue sample, measuring cytokine production to function as a predictive biomarker of clinical response^25^. Mice bearing MOC2 were treated with either dual ICI alone or 10 Gy mean tumor dose from ^177^Lu-NM600 + dual ICI (3 doses pre- and post-RPT treatment). At day 12 after RPT, MOC2 tumors were harvested and core needle biopsies were collected. Core needle biopsies were cut using the an automated instrument called elive^TM^ *Edge* and encapsulated in elive gel for *ex vivo* culture. Encapsulated cores were treated sequentially with control IgG for 24 hours followed by dual PD-L1 and CTLA-4 blockade for an additional 24 hours. Supernatants were collected at 4 hours, 24 hours, and 48 hours were analyzed via multiplex cytokine assay to quantify cytokine dynamics over time.

**Figure 2.**
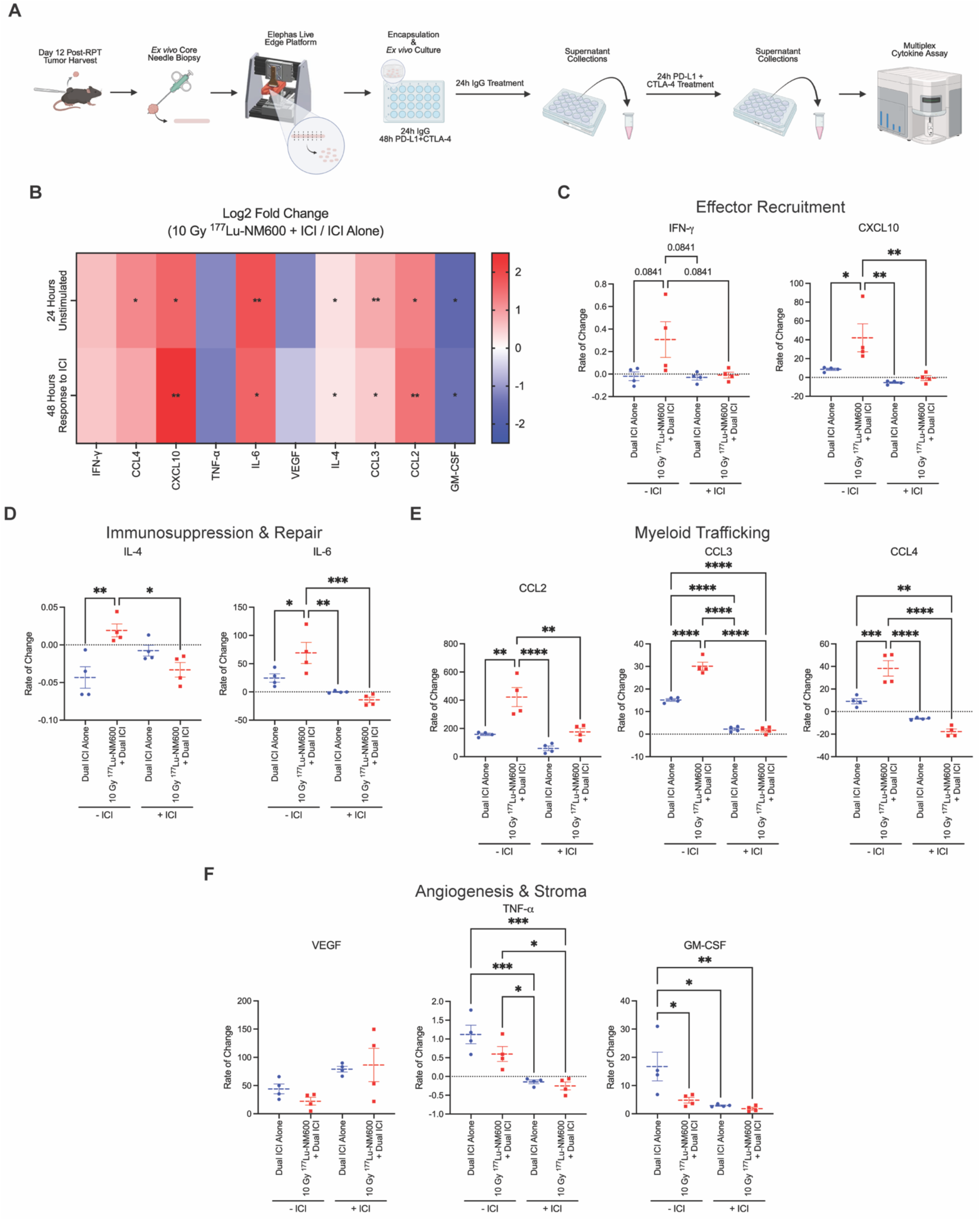
Dynamic inflammatory and chemokine landscapes driven by combination ^177^Lu-NM600 and dual immune checkpoint inhibition. A-K) MOC2 tumor-bearing mice were treated with dual ICI or 10 Gy ^177^Lu-NM600 + dual ICI. Dual ICI was administered day −6, −3, 0, 3, 6, and 9 with ^177^Lu-NM600 administered on day 1. Tumors were harvested at day 12 and core needle biopsies were taken *ex vivo* and cultured using the Elephas Live Edge workflow. A) Schematic for Elephas workflow. B) Heat map of Log2 fold change analysis (10 Gy ^177^Lu-NM600 + ICI/ ICI alone) of 10 secreted cytokines and chemokines following 24-hour IgG *ex vivo* treatment and 24-hour dual ICI *ex vivo* treatment. C) Analysis of IFN-γ and CXCL10 rate of change following 24-hour IgG *ex vivo* treatment and 24-hour dual ICI *ex vivo* treatment. D) Analysis of IL-4 and IL-6 rate of change following 24-hour IgG *ex vivo* treatment and 24-hour dual ICI *ex vivo* treatment. E) Analysis of CCL2, CCL3, and CCL4 rate of change following 24-hour IgG *ex vivo* treatment and 24-hour dual ICI *ex vivo* treatment. F) Analysis of VEGF, GM-CSF, and TNF-α rate of change following 24-hour IgG *ex vivo* treatment and 24-hour dual ICI *ex vivo* treatment. A-G) N=4 all groups. *p < 0.05, **p < 0.01, ***p < 0.001, ****p < 0.0001 by one sample T-test (B). *p < 0.05, **p < 0.01, ***p < 0.001, ****p < 0.0001 by one-way ANOVA (C-F).

To identify secretome changes between tumors from mice treated with an optimized heterogeneous dose of RPT versus those that did not, we calculated the fold change in cytokines and chemokines secreted by these tumors *ex vivo* after 24 hours in culture in the absence of ICI and after an additional 24 hours in culture during which the samples were cultured with dual ICI (Figure 2B). In the absence of ICI in culture, tumors from mice treated with both RPT and ICI had significantly elevated CXCL10, IL-4, IL-6, CCL2, CCL3, and CCL4 compared to tumors from mice that were treated with the dual ICI. Among the pro-inflammatory cytokines, only GM-CSF decreased with the addition of RPT compared to ICI alone, and IFN-γ, VEGF, and TNF-α remained unchanged. When anti-CTLA-4 and anti-PD-L1 were added to the cultured tumor for 24 hours, this pattern was retained (Figure 2B).

Despite this similar secretome profile at both time points, the kinetics of different cytokine levels shifted significantly during the second 24 hours in the presence of ICI. We calculated the rate of change of cytokine secretion between 4 hours and 24 hours when no additional ICI was added in to the *ex vivo* tumors (Figure 2C-F and S1) and the rate of change between 24 and 48 hours when ICIs were present in the culture (Figure 2C-F and S1). For most cytokines accumulation over time was more rapid in the first 24 hours of culture for tumors derived from mice receiving ^177^Lu-NM600 + dual ICI, with the exception being GM-CSF. Interestingly, this rapid increase declined across most chemokines and cytokines measured during the second 24-hour period, where dual ICIs were present, with notable reduced production of IL-4, IL-6, and CCL4. Prior live tumor fragment studies have associated increased cytokine production following *ex vivo* checkpoint blockade, including induction of IFN-γ and CXCL10, with ICI-responsive tumors^24,25^. In contrast, tumors preciously treated with RPT in our study exhibited a chemokine-rich baseline before additional *ex vivo* checkpoint blockade, followed by reduced accumulation of several cytokines during subsequent ICI exposure. This distinct kinetic pattern suggests that prior RPT alters the baseline immune state from which tumors respond to additional checkpoint inhibition. Together, these results suggest that RPT functionally reprograms the TME by establishing a chemokine-rich baseline and altering the kinetics of subsequent cytokine responses to dual ICI.

### Optimized RPT dose enhances dendritic cell activation and migration

To delineate the immunological mechanism underlying the antitumor efficacy of RPT dose heterogeneity in combination with ICIs, we investigated whether the combination therapy augmented DC subtype frequencies and activation in the TDLN at 1 or 2 weeks after RPT injection. Because cDC1 activation is an essential first step required for tumor antigen cross-presentation and co-stimulatory signaling to naïve CD8 T cell, we evaluated cDC1 subtype frequencies and activation by flow cytometry. We distinguished between migratory tissue cDC1s (CD103+), which actively traffic antigen from the tumor to the TDLN, and the broader lineage-defined cDC1s (XCR1+), which include lymph node resident populations^26,27^. An immunologically optimized dose of ^177^Lu-NM600 in combination with dual ICI led to significant increase in the number and activation of migratory, CD103+ cDC1s in the TDLN compared to either RPT or dual ICI alone in MOC2-bearing mice at day 7 (p<0.05; Figure 3A). This was accompanied by elevated cell surface expression of the activation markers CD80 and MHCII (p<0.001 and p<0.05; Figure 3B). However, the increased levels of CD80 and MHCII on CD103+ cDC1s was not observed at day 12 (Figure 3C). Combination treatment also significantly increased expression of CD80 and MHCII within the broader, XCR1+ cDC1 population (p<0.01 and p<0.05; Figure 3D). This increased CD80 and MHCII expression on XCR1+ cDC1s was not sustained to day 12 (Figure 3E). In KP sarcoma-bearing mice, combination therapy significantly increased cDC1 infiltration (p<0.001; Figure 3F) as well as CD80 and MHCII MFI at day 7 (CD80: p<0.05 and MHCII: p<0.01; Figure 3G). The increased CD80 and MHCII MFI within this cDC1 population persisted through two weeks after RPT injection (p<0.05; Figure 3H).

**Figure 3.**
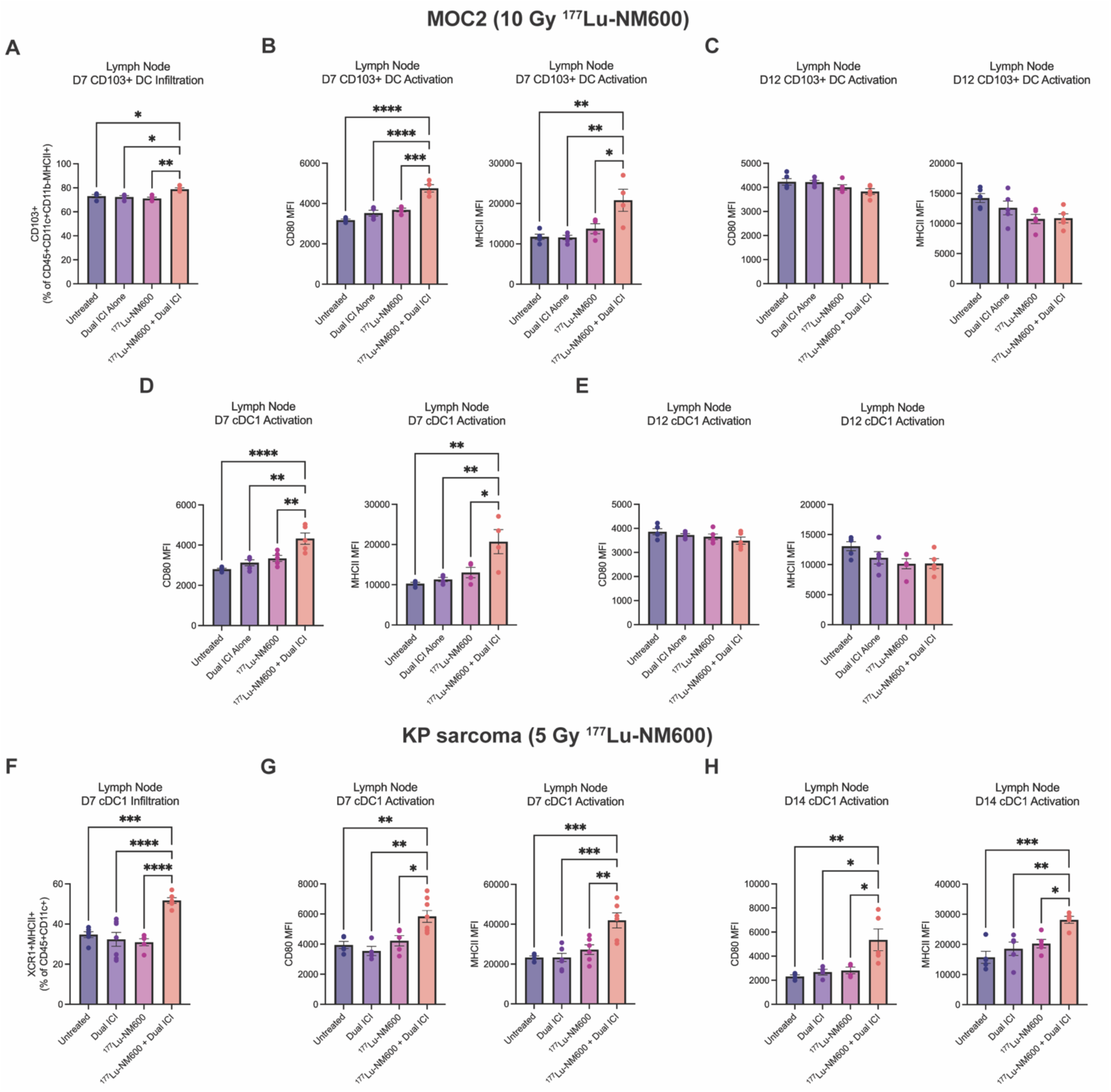
Radiation dose heterogeneity from ^177^Lu-NM600 in combination with dual immune checkpoint inhibition leads to infiltration and activation of CD8-priming dendritic cells in the tumor draining lymph node. A-E, I-J) MOC2 tumor-bearing mice were treated with no treatment, dual ICI (anti-PD-L1 + anti-CTLA-4), 10 Gy ^177^Lu-NM600, or 10 Gy ^177^Lu-NM600 + dual ICI. Dual ICI was administered on day −3, 0, 3 with ^177^Lu-NM600 administered on day 1. Tumors and tumor draining lymph nodes (TDLN) were harvested at day 7 and 12. A) Analysis of migratory dendritic cells, CD103+ (CD45+CD11b-CD11c+MHCII+CD103+) infiltration in the TDLN at day 7. B) Analysis of CD80 and MHCII MFI of CD103+ dendritic cells in the TDLN at day 7. C) Analysis of CD80 and MHCII MFI of CD103+ dendritic cells in the TDLN at day 12. D) Analysis of CD80 and MHCII MFI of conventional type I dendritic cells, cDC1 (CD45+CD11b-CD11c+MHCII+XCR1+) in the TDLN at day 7. E) Analysis of CD80 and MHCII MFI of cDC1 in the TDLN at day 12. F-H, K-L) KP sarcoma tumor-bearing mice were treated with no treatment, dual ICI (anti-PD-L1 + anti-CTLA-4), 5 Gy ^177^Lu-NM600, or 5 Gy ^177^Lu-NM600 + dual ICI. Dual ICI was administered on day −3, 0, 3 with ^177^Lu-NM600 administered on day 1. Tumors and tumor draining lymph nodes (TDLN) were harvested at day 7 and 14. F) Analysis of cDC1 infiltration in the TDLN at day 7. G) Analysis of CD80 and MHCII MFI of cDC1 in the TDLN at day 7. H) Analysis of CD80 and MHCII MFI of cDC1 in the TDLN at day 12. I) Analysis of CD8+ T cells (CD45+CD3+ CD4-CD8+) infiltration at day 14. J) Analysis of CD69 and IFN-γ MFI of CD8+ T cells in the tumor at day 12. K) Analysis of CD8+ T cells (CD45+CD3+ CD4-CD8+) infiltration at day 14. L) Analysis of CD69 and IFN-γ MFI of CD8+ T cells in the tumor at day 14. A-L) N=4 all groups. *p < 0.05, **p < 0.01, ***p < 0.001, ****p < 0.0001 by one-way ANOVA (A-L).

### Spatial transcriptomics reveals distinct heterogeneous tumor microenvironments and treatment-associated immune remodeling

While flow cytometry and cytokine analysis establishes effects of RPT+ICI combination in both the tumor and TDLN, capturing localized responses driven by RPT dose heterogeneity requires spatially resolved analyses. We performed spatial transcriptomics to map regional transcriptomic heterogeneity across TMEs following ^177^Lu-NM600 and dual ICI treatment in mice bearing MOC2 tumors. Gene expression profiles were measured by Visium HD spatial transcriptomics. Following quality-control filtering (Figure S2), a balanced reference was constructed across all samples, and Principal Component Analysis (PCA) followed by shared-nearest-neighbor graph clustering identified 15 distinct transcriptional states, which were we visualized by Uniform Manifold Approximation and Projection (UMAP) (Figure 4A-3E). The 15 clusters were identified as *Pthlh* (parathyroid hormone-like hormone)/*Slpi* (secretory leukocyte peptidase inhibitor) epithelial, cycling stromal/vascular, proliferative epithelial, *Postn*+ (periostin) cancer associated fibroblasts, neural-like, hypoxic/stress, inflammatory polymorphonuclear (PMN)-like myeloid, interferon (IFN)-high antigen-presenting (APC) myeloid, *Aqp5* (aquaporin 5)/*Ncam1* (neural cell adhesion molecule 1)-expressing, endothelial, cytotoxic T/NK, mast cells, osteoclast-like myeloid, *Pi16* (peptidase inhibitor 16)/*Cxcl12* (C-X-C motif chemokine ligand 12) stroma, skeletal muscle (Figure 4C and 4D). All clusters were present in each sample, but their relative frequency differed between the ICI-only treated tumor and the three ^177^Lu-NM600 + dual ICI treated tumors (Figure 4E). To assess how these transcriptional states were spatially organized within the intact tissue architecture, cluster identities were mapped directly onto the spatial coordinates of each tissue section (Figure 4F). Spatial projection demonstrated distinct regional compartmentalization across the TME, with defined epithelial and stromal niches surrounding localized areas of immune infiltration (Figure 4F). Quantitative evaluation of cluster proportions across the QC-retained 8-μm bins demonstrated substantial differences in microenvironmental composition across specimens (Figure 4G).

**Figure 4.**
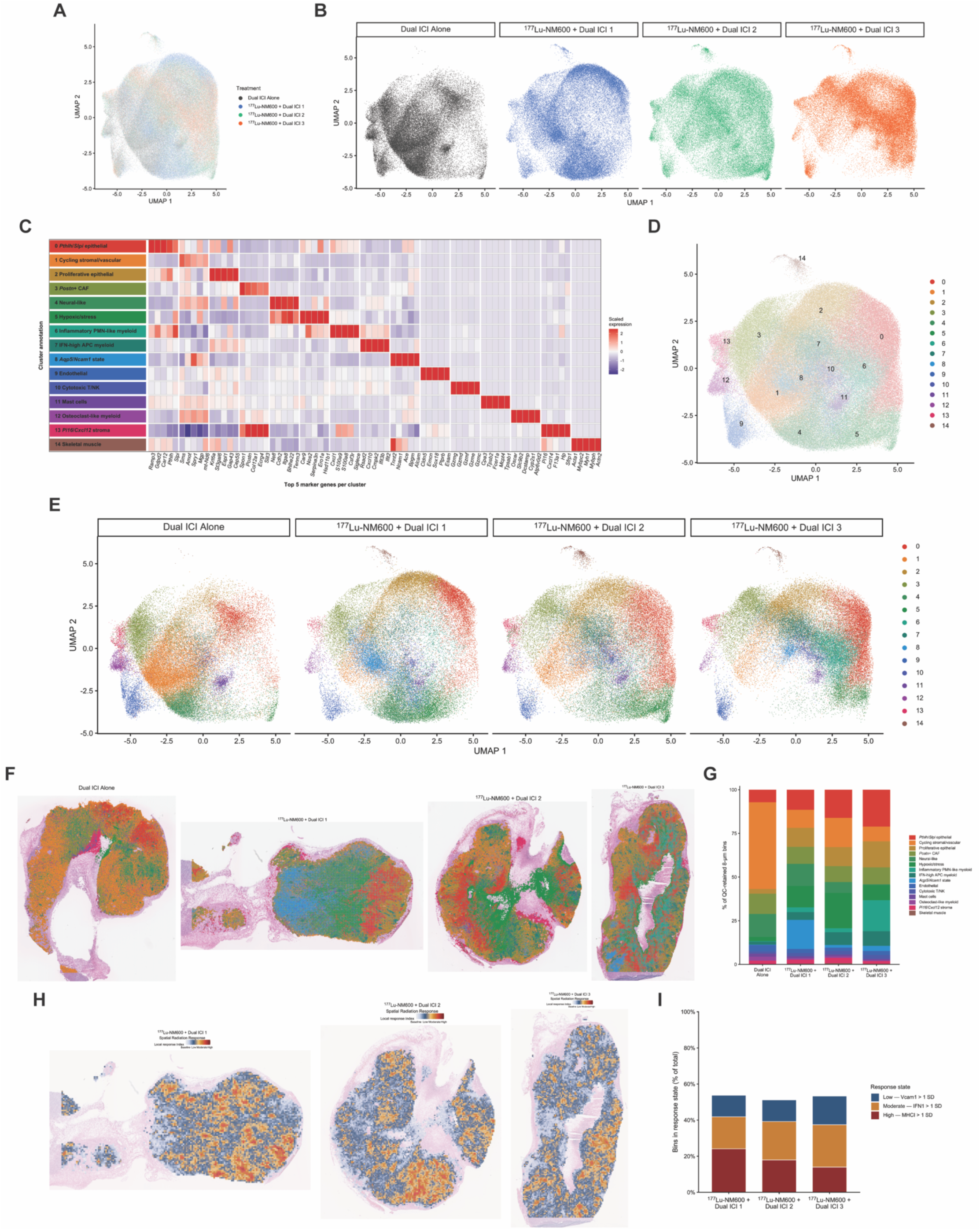
Spatial transcriptomics maps tumor architecture and heterogeneous radiation-associated response states. A-I) MOC2 tumors were collected 7 days following either dual ICI alone (1 tumor) or 10 Gy ^177^Lu-NM600 + dual ICI (3 tumors). A) UMAP visualization of the balanced spatial transcriptomic reference colored by specimen. B) UMAP visualization separated by specimen. C) Heatmap of tup five marker genes for each of 15 transcriptional clusters. D) UMAP visualization of the 15 transcriptional clusters identified as *Pthlh*/*Slpi* epithelial, cycling stromal/vascular, proliferative epithelial, *Postn*+ cancer-associated fibroblast, neural-like, hypoxic/stress, inflammatory PMN-like myeloid, *Ifn*-high APC/myeloid, *Pi16*/*Cxcl12* stroma, and skeletal muscle. E) UMAP visualization of transcriptional clusters separated by specimen. F) Spatial projection of transcriptional cluster identities onto the full-resolution tissue sections. G) Relative abundance of transcriptional clusters across QC-retained 8-μm bins for each specimen. H) Fixed 64-μm spatial response landscapes generated from VCAM1-, type I interferon (IFN1)-, and MHCI-associated transcriptional response programs standardized relative to the ICI specimen. I) Percentage of total QC-retained tissue bins occupying Low (VCAM1-associated), Moderate (IFN1-associated), or High (MHCI-associated) response states.

To evaluate local tissue responses in the absence of direct physical dose co-registration on histological sections, we examined radiation-dose-dependent transcriptional programs as indicators of local biological response rather than direct measures of absorbed radiation dose. Specifically, pathway regulating endothelial adhesion (*Vcam1*)^12,17^, type I interferon signaling (IFN1)^12,15,16^, and antigen presentation (MHC1)^12,13^ have been reported to exhibit differential induction across radiation doses, including with RPT^10^. Using ICI as a baseline reference, response scores were standardized to the ICI specimen, and individual 8-μm bins were considered positive for a given response program when the corresponding score exceeded 1 SD above the ICI reference mean. To assign mutually exclusive functional states to bins expressing multiple signatures, we applied a hierarchical priority scheme (MHC1 > IFN1 > Vcam1), yielding, Baseline, Low (Vcam1-high), Moderate (IFN1-high), and High (MHCI-high) composite response states. These classifications were aggregated into fixed 64-μm spatial neighborhoods to visualize regional biological response across each tissue section. Spatial mapping of these classified bins revealed heterogeneous response landscapes across treated tumors at both native 8-μm and aggregated 64-μm resolutions (Figure S3A and Figure 4H). Quantitative profiling demonstrated a greater proportion of response-active tissue in each of the three ^177^Lu-NM600 + dual ICI specimens relative to the ICI specimen, with increased representation of IFN1- and MHCI-associated response states (Figure 4I). Because the hierarchical classification is intentionally mutually exclusive, we additionally analyzed the VCAM, IFN1, and MHCI axes independently to determine the extent to which these response programs co-occurred. Pairwise tissue-state analyses demonstrated that these three programs were partially overlapping but nonidentical, with substantial tissue occupying single-program or discordant response states and a smaller fraction occupying joint high-response states (Figure S3B). These findings reveal a TME that does not exist as a single uniform transcriptional state but rather as a heterogeneous response landscape in which gradients of radiation-dose-response programs can exist in spatially restricted patterns that may reflect dose distribution when tissue type and hypoxia are accounted.

### Migration-associated DC-like-cell states preferentially localize in regions with greater biological radiation response

We sought to resolve the spatial distribution of immune cell populations within this landscape of dose-dependent spatial transcriptomics data. To achieve a higher-resolution characterization of intratumoral immune dynamics we isolated and re-clustered the immune compartment from our spatial transcriptomics dataset (Figure 5A). Re-clustering resolved seven distinct immune cell lineages, including T cells, neutrophils, monocytes, macrophages, IFN-high antigen-presenting myeloid, osteoclast-like myeloid, and mast cells (Figure 5B and 5C). Evaluation of immune-state composition across specimens demonstrated substantial reorganization of immune cell states, with ^177^Lu-NM600 + dual ICI combination treated tumors showing increased representation of antigen-presenting and interferon-responsive myeloid states relative to the specimen treated with dua ICI-alone (Figure 5D).

**Figure 5.**
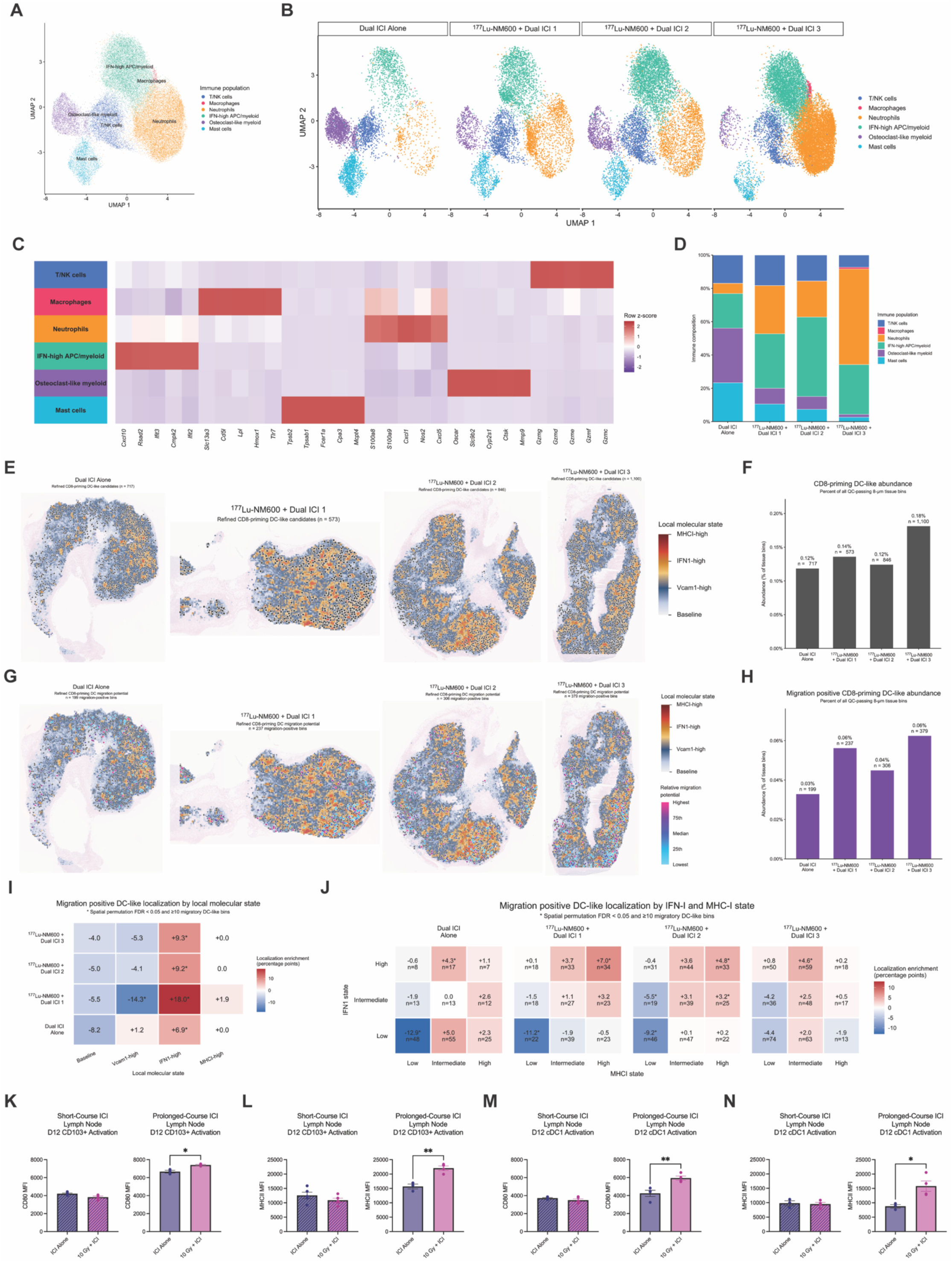
Optimized heterogeneous RPT increases migration positive CD8-priming DC-like cells and promotes their localization to IFN1/MHCI-high tumor states. A-H) MOC2 tumors were collected 7 days following either dual ICI alone (1 tumor) or 10 Gy ^177^Lu-NM600 + dual ICI (3 tumors). A) UMAP visualization of the re-clustered immune compartment identifying T/NK cells, macrophages, neutrophils, IFN-high APC/myeloid, osteoclast-like myeloid, and mast cells. B) UMAP visualization of immune populations separated by specimen. C) Heatmap of representative marker genes defining each immune population. D) Relative composition of immune populations across specimens. E) Spatial distribution of refined CD8-priming DC-like candidate bins across tumor sections. F) Quantification of refined CD8-priming DC-like candidate abundance normalized to all QC-passing 8-μm tissue bins. G) Spatial distribution of migration-positive refined DC-like candidate bins across tumor sections. H) Quantification of migration-positive refined DC-like candidate abundance normalized to all QC-passing 8-μm tissue bins. I) Localization enrichment of migration-positive DC-like candidate bins across Baseline-like, Low-like, Moderate-like, and High-like radiation-associated response states. J) Localization enrichment of migration-positive DC-like candidate bins across joint IFN1 and MHCI response states. K-N) MOC2 tumor-bearing mice were treated with dual ICI (anti-PD-L1 + anti-CTLA-4) or 10 Gy ^177^Lu-NM600 + dual ICI. Dual ICI was administered day −3, 0, 3 with ^177^Lu-NM600 on day 1 (hashed bar graph, data originally presented in Figure 3). Dual ICI was administered day −6, −3, 0, 3, 6, and 9 with ^177^Lu-NM600 administered on day 1 (solid bar graph). Tumor draining lymph nodes (TDLN) were harvested at day 12 post-RPT treatment. K) Analysis of CD80 of CD103+ dendritic cells (CD45+CD11b-CD11c+MHCII+CD103+) in the TDLN at day 12. L) Analysis of MHCII MFI of CD103+ dendritic cells in the TDLN at day 12. M) Analysis of CD80 of conventional type I dendritic cells, cDC1 (CD45+CD11b-CD11c+MHCII+XCR1+) in the TDLN at day 12. N) Analysis of MHCII MFI of CD103+ dendritic cells in the TDLN at day 12. I-J) Positive enrichment indicates preferential localization relative to available tissue; *BH-adjusted spatial permutation FDR < 0.05 with ≥10 candidate bins. K-N) N=4 all groups. *p < 0.05, **p < 0.01, ***p < 0.001, ****p < 0.0001 by unpaired T-test.

Because effective anti-tumor immunity depends on DC activation, antigen presentation, and trafficking to the TDLN, we next examined DC-like transcriptional states within the spatial dataset. Using a stringent transcriptional definition incorporating DC-lineage, MHCII/APC, cDC1/cross-presentation programs, we identified refined CD8-priming DC-like candidate bins and further characterized migration-positive subsets based on expression of migration-associated genes (Figure 5E-H). Spatial projections demonstrated that these DC-like candidate populations occupied discrete regions across tumor sections (Figure 5E and 5G). Quantification showed greater representation of CD8-priming DC-like candidates and migration-positive DC-like candidates in each of the three ^177^Lu-NM600 + dual ICI specimens relative to the specimen treated with dual ICI-alone (Figure 5F and 5H). These spatially resolved candidate populations therefore provided a framework for testing whether migratory DC-like states preferentially localized within specific radiation-associated response microenvironments.

To determine whether migration-positive DC-like candidate bins preferentially occupied particular response states, we first quantified the localization across the hierarchical Baseline, Low, Moderate, and High response landscape. Migration-positive DC-like candidates were diminished among baseline-like regions and enriched within radiation-responding tissue states, with prominent enrichment in IFN1-associated response areas across ^177^Lu-NM600 + dual ICI specimens (Figure 5I). Because the hierarchical classifier assigns each bin to a single dominant response state, we examined migration-positive DC localization across independent pairwise response saxes, including VCAM, IFN1, and MHCI. For each response state, enrichment was calculated as the percentage of migration-positive DC-like candidate bins occupying that state minus the percentage of available tissue represented by the same state. This analysis demonstrated preferential localization of migration-positive DC-like candidates within response-active microenvironments across independent pairwise response axes. Migration-positive DC-like candidates were consistently depleted from regions in which both response axes were low and were enriched as radiation-response activity increased, with particularly prominent localization within IFN1-high regions accompanied by intermediate-to-high MHC1 or VCAM responses (Figure 5J and Figure S4A). This enrichment reflects preferential localization relative to available tissue rather than simple co-occurrence with abundant response states. Together, these analyses indicate that migration-associated DC-like states are nonrandomly distributed across the heterogeneous radiation-response landscape and preferentially occupy immunologically active tissue regions generated following RPT treatment. These transcriptional features are consistent with enhanced migration potential but do not directly demonstrate trafficking to the TDLN.

Based on the greater therapeutic response in adding RPT to a well-established ICI regimen with extended ICI treatment (Figure 1S-U), we evaluated whether improved therapeutic response was due to prolonged DC activation. We previously demonstrated that a short course of dual ICI in combination with 10 Gy ^177^Lu-NM600 led to activation of cDC1s at day 7 but activation was not sustained to day 12 (Figure 3). To determine the effect of prolonged dual ICI treatment on DC activation, we evaluated conventional type I DC (cDC1) activation within the TDLN at day 12 post-RPT. Mice bearing MOC2 were treated with either dual ICI alone or 10 Gy ^177^Lu-NM600 + dual ICI (3 doses pre- and post-RPT treatment). At day 12, 10 Gy ^177^Lu-NM600 with prolonged dual ICI led to increased expression of CD80 MFI (p=0.0211; Figure 5K) and MHCII MFI (p=0.0035; Figure 5L) on migratory DCs compared to dual ICI alone. Additionally, combination treatment with prolonged dual ICI led to increased CD80 MFI (p=0.0087; Figure 5M) and MHCII MFI (p=0.0249; Figure 5N) on lineage-defined cDC1s. These results contrast with the shorter-term ICI treatment in combination with 10 Gy ^177^Lu-NM600 where no increased expression of CD80 or MHCII MFI was detected at day 12.

### Optimized dose heterogeneity from RPT induces expansion of tumor-specific CD8 T cells

Given that DC activation in the TDLN is required to prime naïve T cells, we assessed whether cDC1 infiltration and activation translated to enhanced T cell responses within the tumor^27,28^. Despite the short-lived cDC1 activation in the TDLN, combination therapy produced a consistent increase in intratumoral CD8 T cell infiltration and activation across both tumor models (Figure 6A-D). In MOC2-bearing mice, combination treatment significantly elevated CD8 T cell tumor infiltration (p<0.05; Figure 6A) and increased T cell expression of the activation markers CD69 and IFN-γ (p<0.01; Figure 6B) relative to monotherapies. Similarly, in KP sarcoma-bearing mice, combination therapy significantly increased tumor infiltration of CD8 T cells (p<0.01; Figure 6C) and CD69 and IFN-γ levels on these T cells (CD69: p<0.01 and IFN-γ p<0.0001; Figure 6D). This data demonstrates that, when applying dosimetric metrics for optimizing the dosing of RPT in combination with dual ICI, selected RPT doses consistently activate dendritic cells and induce tumor infiltration and activation of CD8 T cells.

**Figure 6.**
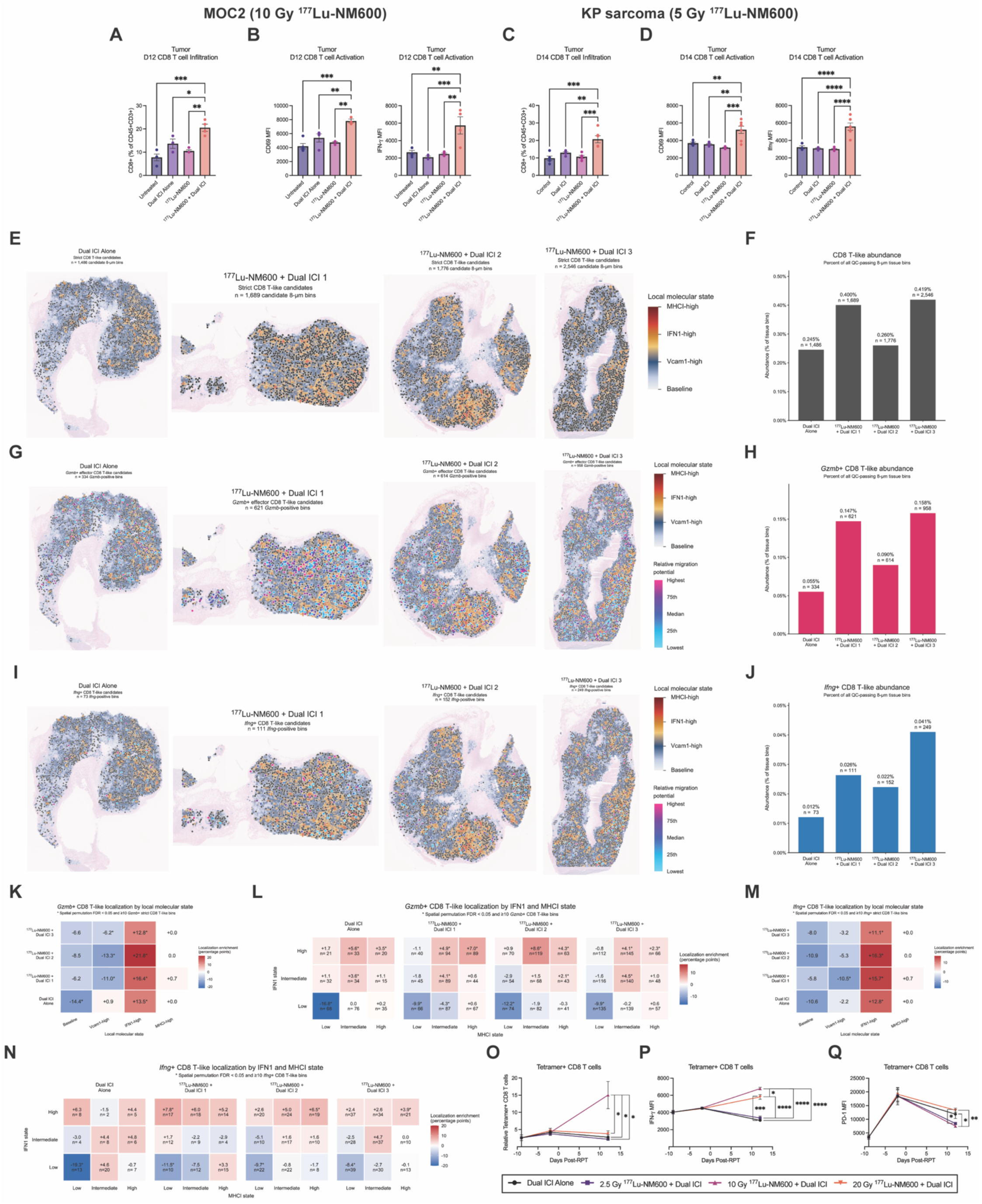
^177^Lu-NM600 in combination with dual ICI promotes spatially organized CD8 T-like effector and antigen-specific responses. A-B) MOC2 tumor-bearing mice were treated with no treatment, dual ICI (anti-PD-L1 + anti-CTLA-4), 10 Gy ^177^Lu-NM600, or 10 Gy ^177^Lu-NM600 + dual ICI. Dual ICI was administered on day −3, 0, 3 with ^177^Lu-NM600 administered on day 1. Tumors were harvested at day 12. A) Analysis of CD8+ T cells (CD45+CD3+ CD4-CD8+) infiltration at day 14. B) Analysis of CD69 and IFN-γ MFI of CD8+ T cells in the tumor at day 12. C-D) KP sarcoma tumor-bearing mice were treated with no treatment, dual ICI (anti-PD-L1 + anti-CTLA-4), 5 Gy ^177^Lu-NM600, or 5 Gy ^177^Lu-NM600 + dual ICI. Dual ICI was administered on day −3, 0, 3 with ^177^Lu-NM600 administered on day 1. Tumors were harvested at day 14. C) Analysis of CD8+ T cells (CD45+CD3+ CD4-CD8+) infiltration at day 14. D) Analysis of CD69 and IFN-γ MFI of CD8+ T cells in the tumor at day 14. E-N) MOC2 tumors were collected 7 days following either dual ICI alone (1 tumor) or 10 Gy ^177^Lu-NM600 + dual ICI (3 tumors). E) Spatial distribution of all strict CD8 T-like candidate bins. F) Quantification of strict CD8 T-like candidate abundance normalized to all QC-passing 8-μm tissue bins. G) Spatial distribution of *Gzmb*-positive strict CD8 T-like candidate bins. H) Quantification of *Gzmb*-positive strict CD8 T-like candidate abundance normalized to all QC-passing 8-μm tissue bins. I) Spatial distribution of *Ifng*-positive strict CD8 T-like candidate bins. J) Quantification of *Ifng*-positive strict CD8 T-like candidate abundance normalized to all QC-passing 8-μm tissue bins. K) Localization enrichment of *Gzmb*-positive strict CD8 T-like candidate bins across Baseline-like, Low-like, Moderate-like, and High-like radiation-associated response states. L) Localization enrichment of *Gzmb*-positive strict CD8 T-like candidate bins across joint IFN1 and MHCI response states. M) Localization enrichment of *Ifng*-positive strict CD8 T-like candidate bins across Baseline-like, Low-like, Moderate-like, and High-like radiation-associated response states. N) Localization enrichment of *Ifng*-positive strict CD8 T-like candidate bins across joint IFN1 and MHCI response states. O-Q) MOC2 tumor-bearing mice were treated with dual ICI or 10 Gy ^177^Lu-NM600 + dual ICI. Dual ICI was administered day −6, −3, 0, 3, 6, and 9 with ^177^Lu-NM600 administered on day 1. O) Analysis of circulating p15E tetramer-positive CD8+ T cells (CD45+CD3+CD4-CD8+Tetramer+) in peripheral blood pre-ICI, pre-RPT, and 12 days post-RPT treatment. P) Analysis of IFN-γ MFI of circulating p15E tetramer-positive CD8+ T cells in peripheral blood pre-ICI, pre-RPT, and 12 days post-RPT treatment. Q) Analysis of PD-1 MFI of circulating p15E tetramer-positive CD8+ T cells in peripheral blood pre-ICI, pre-RPT, and 12 days post-RPT treatment. A-D) N=4 all groups. O-Q) N=3 all groups. *p < 0.05, **p < 0.01, ***p < 0.001, ****p < 0.0001 by one-way ANOVA (A-D). Positive enrichment indicates preferential localization relative to available tissue; *BH-adjusted spatial permutation FDR < 0.05 with ≥10 candidate bin (E, G, I, K). *p < 0.05, **p < 0.01, ***p < 0.001, ****p < 0.0001 by two-way ANOVA Mixed-effects (O-Q).

We evaluated the spatial abundance and effector-state composition of tumor-infiltrating CD8 T-like candidate bins (Figure 6E-J). Conservative CD8 T-like candidates were defined by co-detection of CD8-lineage and T cell receptor-associated transcripts. Spatial mapping revealed greater representation of CD8 T-like candidate bins in the ^177^Lu-NM600 + dual ICI specimens relative to the ICI specimen (Figure 6E), and tissue-normalized quantification showed a corresponding increase in their abundance (Figure 6F). We further examined cytotoxic and cytokine-associated effector states using *Gzmb* and *Ifn-γ* expression. The combination-treated specimens exhibited increased representation of both *Gzmb*-positive and *Ifng*-positive CD8 T-like candidate bins relative to the ICI specimen (Figure 6G-J). Using the same response-state localization framework applied to migration-positive DC-like candidates, both *Gzmb*-positive and *Ifng*-positive CD8 T-like states were depleted from response-low regions and preferentially enriched within response-active regions (Figure 6K-N). *Gzmb*-positive CD8 T-like candidates showed a particularly consistent pattern of depletion from IFN1-low/MHCI-low regions and enrichment within IFN1-high, intermediate-to-high MHCI states (Figure 6K-L), whereas *Ifng*-positive CD8 T-like candidates demonstrated a similar overall bias toward IFN1-high regions with greater variability across individual MHCI states (Figure 6M-N). Comparable localization patterns were observed across the VCAM x IFN1 and VCAM x MHCI response states with low response activity and preferentially enriched within regions exhibiting elevated VCAM together with increased IFN1 and MHCI response states (Figure S4B-C). These finding are consistent with a TME characterized by regions of radiated tumor that exhibit increased accumulation of cytotoxic and effector-associated CD8 T-like states alongside migration-associated DC-like populations.

We previously reported that a heterogeneous radiation dose from RPT promotes CD8 T cell clonal expansion^10^. Like dual ICI alone, however, mice eventually succumbed to their tumors. Given the durable response both in survival (Figure 1S-U) and cDC1 activation (Figure 5K-N) of mice receiving the extended ICI treatment combined with RPT, we next sought to determine if this response translated to an increase in activated, circulating tumor-specific CD8 T cells that also depended on a heterogeneous radiation dose. Using our extended ICI regimen, we treated mice bearing MOC2 with dual ICI, 2.5 Gy ^177^Lu-NM600 + dual ICI, 10 Gy ^177^Lu-NM600 + dual ICI, or 20 Gy ^177^Lu-NM600 + dual ICI. Blood was drawn by submandibular vein prior to ICI treatment, prior to RPT treatment, and 12 days following RPT. To determine the effect on tumor-specific CD8 T cells, we leveraged the fact that MOC2 cells express the known T cell antigen, p15E expression arises from the epigenetic de-repression of an endogenous retrovirus within the derivative cell line^29,30^. This antigen is not expressed in normal C57BL/6 mice^29^. We utilized an H-2Kb-restricted p15E MHCI tetramer to track circulating tumor-specific CD8 T cells over time^31,32^.

Prior to dual ICI treatment and RPT treatment, circulating p15E+ CD8 T cell frequencies remained uniformly low and equivalent across all experimental cohorts, consistent with primary ICI resistance observed in the MOC2 model. At day 12 following RPT treatment, 10 Gy mean tumor dose from ^177^Lu-NM600 led to a significant increase in tetramer^+^ CD8^+^ T cells compared to dual ICI alone or 2.5 Gy or 20 Gy mean tumor dose from ^177^Lu-NM600 combined with ICIs (p<0.05; Figure 6O). To characterize this expansion of tumor-specific CD8 T cells, we quantified IFN-γ in circulating tetramer^+^ CD8 T cells. The optimized heterogeneous dose of 10 Gy ^177^Lu-NM600 in this model led to a significant increase in IFN-γ expression among tetramer^+^ CD8^+^ T cells as compared to ICI alone or ICI combined with 2.5 Gy or 20 Gy mean tumor dose from ^177^Lu-NM600 (p<0.001; Figure 6P). While 2.5 Gy mean tumor dose in this model failed to increase IFN-γ on these T cells compared to ICI alone, 20 Gy produced intermediate IFN-γ amounts, indicating potential for effectiveness that was constrained by concurrent induction of lymphopenia (Figure 1P). Surface PD-1 exhibited a distinct pattern, with the highest expression observed on tetramer^+^ CD8^+^ T cells following dual ICI alone or combination of 20 Gy mean tumor dose from ^177^Lu-NM600 with dual ICI, whereas combination with 10 Gy and 2.5 Gy mean tumor dose from ^177^Lu-NM600 maintained significantly lower levels (p<0.05; Figure 6Q). Crucially, only the 10 Gy mean tumor dose from ^177^Lu-NM600 achieved higher IFN-γ MFI while maintaining low PD-1 levels following treatment of MOC2 tumors. In contrast, the elevated PD-1 in the 20 Gy mean tumor dose combination coincided with reduced IFN-γ relative to 10 Gy mean tumor dose. These data demonstrate that adding appropriately dosed RPT to ICI treatment can overcome primary ICI resistance, driving durable local tumor control and systemic expansion and activation of tumor-specific CD8^+^ T cells.

## DISCUSSION

Our results suggest that dose optimization of RPT may be critically important in translational studies that seek to promote adaptive anti-tumor immunity through combination with ICI therapy. Our findings also support the potential validity of our hypothesized dose optimization metrics for identifying effective administered activities of RPT for combining with ICIs for a given drug, tumor type, and host. Our findings demonstrate that combining an immunologically optimized, heterogeneous dose of ^177^Lu-NM600 with dual ICIs helps overcomes primary checkpoint resistance and enhances treatment sensitivity across poorly immunogenic tumors.

Building on our prior finding that an immunologically optimized RPT dose stimulates anti-tumor immunity as a monotherapy, this study indicates that integrating this dosing strategy with dual checkpoint blockade produces enhanced local and systemic anti-tumor responses. Mechanistically, an immunologically optimized, heterogeneous dose of ^177^Lu-NM600 elicits an in situ vaccination effect, promoting cDC1 trafficking and priming in the TDLNs to expand functional, tumor-antigen-specific CD8 T cells. Spatial transcriptomic profiling further revealed regionally heterogeneous radiation-associated response niches characterized by IFN1, MHCI, and VCAM1-associated programs, with preferential localization of migratory DC-like and CD8 T-like states within response-active regions. Importantly, the optimal RPT dose could be predicted from spatial dose-distribution characteristics rather than therapeutic response alone, suggesting intratumoral dose heterogeneity may provide a marker for immunologically informed RPT dosing. Furthermore, incorporating RPT treatment into an ongoing ICI regimen and maintaining checkpoint blockade after RPT promotes this cDC1 activation state, ultimately reshaping the tumor secretome into an effector-recruiting profile while restraining pro-inflammatory and angiogenic overdrive.

In mixed B16-mFAP^+^/mFAP^−^ tumors, image-based dosimetry predicted 5 Gy ^177^Lu-PNT6555 as the dose that simultaneously preserved low-dose regions and achieved high-dose regions, and this dose subsequently produced the greatest therapeutic benefit when combined with dual ICI. Applying the same spatial dosimetric framework to the distinct MOC2 model treated with ^177^Lu-NM600 identified 10 Gy as the predicted immunologically optimized dose, which was subsequently associated with maximal cDC1 activation and improved therapeutic efficacy. Moreover, an independently established immunologically optimized dose of 5 Gy ^177^Lu-NM600 enhanced ICI efficacy in KP sarcoma, supporting the generalizability of this dosing strategy across preclinical tumor types.

Optimizing the immunogenic potential of RPT requires moving beyond conventional radiobiological paradigms centered on delivering a maximal tolerated injected activity with the intent of maximizing induction of DNA damage in tumor cells from radiation. Our findings suggest that the intratumoral distribution of absorbed dose is a critical driver of anti-tumor immunity following RPT. We therefore propose a predictive framework for using dosimetry to optimize RPT dosing in combination with ICIs. Our dosing approach is defined not by a single mean absorbed dose, but by a therapeutic window in which sufficient high-dose regions in the TME are retained to promote tumor immunogenicity while low-dose regions in the TME and systemic immune competence are preserved. At higher doses, loss of low-dose regions and systemic lymphopenia coincide with diminished immune activation, whereas insufficient dosing fails to generate the same immunogenic response. Collectively, our findings support image-based spatially-resolved dosimetry as potentially predictive radiographic marker for selecting RPT doses likely to cooperate with and overcome resistance to ICIs. This may be most feasible for RPT with a theranostic pair of isotopes to enable personalized dosimetry in advance of therapy.

Spatial transcriptomic profiling demonstrates that our heterogeneous dosing strategy was accompanied by marked regional heterogeneity in immuno-radiobiological response. ^177^Lu-NM600 + dual ICI generated spatially distinct response-active regions characterized by VCAM1-, IFN1-, MHCI-associated programs rather than a uniform tumor-wide transcriptional response. Migratory DC-like and effector CD8 T-like states preferentially localized within these response-active microenvironments, providing a tissue-level link between regional radiation-associated responses and the broader cDC1-CD8 T cell immune axis observed by flow cytometry. These finding support a model in which heterogeneous RPT dosing produces a spectrum of immunologically distinct local microenvironments that collectively facilitate antigen presentation, immune recruitment, and cytotoxic activity.

Mechanistically, our findings identify cDC1 activation within the TDLN as a central component of the systemic anti-tumor immune response generated by combination RPT and dual ICI. Following ^177^Lu-NM600 treatment, migratory tissue-derived cDC1s and lineage-defined cDC1 populations exhibited increased infiltration and upregulated co-stimulatory markers within the TDLN. Crucially, temporal evaluation revealed a pivotal therapeutic nuance. While short-course ICI combined with RPT initiated cDC1 priming, this activation subsided. However, extending duration of ICI administration post-RPT sustained cDC1 co-stimulatory marker expression. The prolonged cDC1 activation coincided with expansion of antigen-specific CD8 T cells while maintaining lower surface PD-1 expression, suggesting that optimal RPT dosing supports T cell activation while limiting the rapid induction of exhaustion observed under higher doses (20 Gy) or dual ICI alone.

To evaluate how these systemic immune responses alter the local TME, we utilized live tumor fragments to profile dynamic secretome kinetics in intact tumor architecture. Prior to secondary *ex vivo* checkpoint challenge, RPT-pretreated tumors exhibited an immune-active secretory profile characterized by increased myeloid and cytotoxic-cell recruitment signals together with suppression of selected pro-tumorigenic growth factors. Upon secondary ex vivo checkpoint challenge, RPT-pretreated tissue sustained these essential effector-recruiting axes by maintaining elevated chemoattractant kinetics, while selectively dampening hyper-inflammatory and pro-angiogenic drivers. Together, these live tissue secretome dynamics indicate that RPT conditions the tumor microenvironment to support enhanced ICI responsiveness by building a sustained recruitment network while preventing inflammatory excess.

We acknowledge several limitations to our study. While syngeneic murine tumor models provide an immunocompetent setting essential for evaluating combination immunotherapy, they do not fully recapitulate the human tumorigenesis, the tumor-immune microenvironment, and metastatic potential. Second, this study evaluated local and systemic responses predominantly in primary tumors rather than spontaneous metastatic disease models. Establishing large primary tumor volumes was necessary to evaluate spatially resolved radiopharmaceutical therapy and preserve low-dose regions, which are more challenging to resolve and interrogate spatially in microscopic lesions. However, this focus on primary lesions restricts direct conclusions regarding treatment efficacy in disseminated, multi-organ metastatic settings. In addition, spatial transcriptomic response states were not co-registered with quantitative intratumoral dosimetry and therefore represent regional biological responses associated with therapy rather than direct measurements of local absorbed radiation dose. Because of cost and resource limitations, spatial transcriptomics analyses are limited to one ICI treated tumors and three ^177^Lu-NM600 + dual ICI treated tumors; consequently, specimen-level spatial differences were interpreted descriptively and require validation in larger cohorts. Nevertheless, establishing this spatial radiobiological framework in intact tumors provides a vital proof-of-concept, demonstrating that optimized RPT dosing can safely sensitize poorly immunogenic, ICI-resistant tumors to ICI therapy and this may guide rational design of clinical trials combining RPT and ICI.

Primary resistance to ICIs has traditionally been viewed as a therapeutic ceiling, while RPT in combination regimens has conventionally been guided by maximum tolerated activity^18,19^. Our findings instead support an immunologically informed dosing paradigm in which image-based spatial dose heterogeneity can be used to identify RPT doses that are most likely to synergize with ICIs. Delivering sufficient high-dose regions while preserving intratumoral low-dose regions can resensitize the TME without triggering systemic immune depletion and generates spatially heterogeneous response niches enriched for antigen-presentation and effector-associated immune programs. Crucially, continuing checkpoint blockade after RPT sustains cDC1 activation within tumor draining lymph nodes, expands systemic antigen-specific CD8^+^ T cells, and maintains an effector-recruiting TME secretome. Together, these findings establish a framework in which the spatial distribution of RPT dose can inform RPT dosing and sequencing with ICIs. Leveraging RPTs to overcome ICI resistance may provide a viable strategy to enhance survival in patients with metastatic cancer.

## METHODS

### Cell lines and Cell culture

The murine KP sarcoma cell line, derived from the transgenic KP sarcoma models as previously described^33^, was obtained from Dr. David Kirsch (University Health Network Princess Margaret Cancer Centre). The murine head and neck squamous cell carcinoma MOC2 cell line was obtained from Dr. Ravindra Upalluri (Brigham and Women’s Hospital and DanaFarber Cancer Institute). B16-F10 cell lines were grown in RPMI supplemented with 10% FBS, 100 U/mL penicillin, and 100 μg/mL streptomycin. KP sarcoma cells were grown in DMEM supplemented with 10% FBS, 100 U/mL penicillin, and 100 μg/mL streptomycin. MOC2 cells were grown in DMEM / F12 at a 2:1 mixture and supplemented with 5% FBS, 100 U/mL penicillin, 100 μg/mL streptomycin, 5 ng/mL epidermal growth factor, 400 ng/mL hydrocortisone, and 5 μg/mL insulin. Cells were incubated in a humidified incubator at 37°C with 5% CO_2_. Cell line authentication was performed per ATCC guidelines using morphology, growth curves, and Mycoplasma testing within 6 months of use.

### Murine tumor models

Mice were housed and treated under an Institutional Animal Care and Use Committee (IACUC) approved protocol (protocol number M005670) at the University of Wisconsin – Madison. Female SvJ mice were purchased at age 6 to 8 weeks from Jackson Laboratory. Female C57BL/6 mice were purchased at age 6 to 8 weeks from Taconic. B16-F10 tumors were engrafted by subcutaneous flank injection of 2 x 10^5^ tumor cells. KP sarcoma tumors were engrafted by orthotopic gastrocnemius injection of 5 x 10^4^ tumor cells. MOC2 tumors were engrafted by subcutaneous flank injection of 2 x 10^6^ tumor cells. Tumor size was determined using digital calipers and volume approximated as (length x width^2^)/2. Mice were randomized immediately before treatment by stratified randomization by tumor size. Only mice with palpable tumors the day before treatment began were included in the study. Treatment began when tumors were well-established, approximately 9 and 7 days after tumor implantation for KP sarcoma ((∼30-50 mm^3^) and MOC2 (∼50-70 mm^3^), respectively. The day of RPT was defined as “day 1” of treatment. Mice were euthanized when tumor size exceeded 20 mm in longest dimension or recommended by an independent animal health monitor for morbidity of moribund behavior. Mice bearing KP sarcoma tumors were monitored for cancer progression morbidity behavior to determine endpoint. Due to the institutional radiation safety protocols and the mice being treated with radioactive isotopes, all investigators were aware of mouse treatment groups. The number of mice in therapy groups was based on expected outcomes and an effort to power experimental design to anticipated effect size, determined in collaboration with biostatisticians, and therefore may vary between treatment groups and tumor models. Mouse therapy experiments were repeated in duplicate. Final replicates are presented for tumor response and aggregate data for survival; number of animals per group is indicated in figure legends.

### Radionuclides and Radiochemistry

^177^Lu was purchased as ^177^LuCl_3_ from SHINE Technologies. Radiochemistry was performed as previously described^20,21^.

### iQID Imaging

Mice bearing subcutaneous MOC2 (C57BL/6) were administered 256 μCi ^77^Lu-NM600 in the lateral tail vein. Mice were euthanized at 24h post-injection and tumors were harvested, embedded in optimal cutting temperature gel and frozen on dry ice. The tumor containing OCT blocks were cut at −16°C at a 20 μm thickness. The slide was placed on the iQID camera and imaged for 2h. iQID scans were acquired for the sections as count rate images, which were subsequently converted into activity concentration maps (Bq/mL) using a previously established calibration curve.

### *Ex vivo* microscale dosimetry estimation

For microscale (∼μm) *ex vivo* dosimetry, the absorbed dose rate of tumor sections at the time of harvesting was estimated using ionizing-radiation Quantum Imaging Detector (iQID) activity image-based Monte Carlo simulations in a tissue-equivalent medium (liquid water). This calculation considered only the self-dose within the individual tumor section, excluding cross-dose contributions from adjacent sections or organs. To calculate the mean total absorbed dose, this single time point dose rate was integrated by applying the specific pharmacokinetic parameters from the corresponding tumor model’s serial *in vivo* SPECT/CT data.

### Radiation therapy (RT)

^177^Lu-NM600 therapy was administered via intravenous tail vein injection on treatment day 0 for KP sarcoma and MOC2 tumor-bearing mice. Table 1 shows injected activity for each RPT dose studied.

**Table 1.** Injected RPT activity for tumor dose prescriptions.

| Tumor Model | RPT Agent | Tumor Absorbed Dose Prescription (Gy) | Injected Activity (μCi) |
| --- | --- | --- | --- |
| B16 | <sup>177</sup>Lu-PNT6555 | 2.5 | 167 |
|  | <sup>177</sup>Lu-PNT6555 | 5 | 350 |
|  | <sup>177</sup>Lu-PNT6555 | 10 | 667 |
|  | <sup>177</sup>Lu-PNT6555 | 20 | 1333 |
| KP sarcoma | <sup>177</sup>Lu-NM600 | 5 | 251 |
| MOC2 | <sup>177</sup>Lu-NM600 | 2.5 | 64 |
|  | <sup>177</sup>Lu-NM600 | 10 | 256 |
|  | <sup>177</sup>Lu-NM600 | 20 | 507 |

**Table 2.** Cancer progression morbidity metrics for KP sarcoma.

| Grade | Description |
| --- | --- |
| 1 | Tumor present on right hind leg with full range of motion |
| 2 | Tumor bearing hind leg extended with maintained leg and paw mobility |
| 3 | Tumor bearing hind leg fully extended with movement in the paw |
| 4 | Complete loss of mobility of tumor bearing hind leg |

### Lymphopenia Assessment

To evaluate ^177^Lu-NM600 effect on lymphopenia, complete blood count (CBC) studies were performed in therapy mice. Groups of MOC2 tumor bearing mice (n=3/group) were treated with 2.5, 10, and 15 Gy from ^177^Lu-NM600 intravenously. Submandibular blood collection was performed on three mice 12 post-injection. CBC analysis was completed using whole blood and an Abaxis VetScan HM5 hematology analyzer (Union City, CA).

### KP sarcoma survival assessment

These were defined prior to the start of the study. Grade 1 was defined as tumor-engrafted with full range of motion and use of the tumor bearing hind leg, Grade 2 was defined as a slight (<90°) extension of tumor bearing hind leg, Grade 3 was defined as a complete extension of tumor bearing hind leg and retained foot mobility, and Grade 4 was defined as complete loss of all mobility of tumor bearing hind leg.

### *Ex vivo* cytokine analysis with Elephas Edge

For *ex vivo* tissue culture tumors were harvested 12 days following RPT treatment^24^. Three core needle biopsies (CNBs) were collected using an 18-guage biopsy gun. CNBs were embedded in agarose and cut using the Elephas Live Edge machine into 1 mm fragments. Live tumor fragments were plated in a 24-well culture plate (∼10 fragments/well). Hydrogel (Elephas eLive Protect) was added to each well and polymerized with UV light. After polymerization, 500 μL of culture media with treatment (IgG or αPD-L1/αCTLA-4) was added to each well. The IgG isotype control and immune checkpoint inhibitor antibodies were added at a concentration of 50 μL/mL. Supernatants were collected for downstream cytokine analysis at 4, 22, 46, and 70 hours after start of culture. Cytokine analysis was performed on the conditioned media collected from the *ex vivo* culture using the BioLegend LegendPlex Mouse Cytokine Release Syndrome Panel.

### Flow cytometry

Flow cytometry was performed as previously described^34^, using fluorescent beans (UltraComp Beads eBeads, 176 Invitrogen) to determine compensation, and fluorescence minus once (FMO) methodology to determine gating. Rainbow beads (Spherotech) were used to set voltages for each flow cytometry timepoint. For *in vivo* analysis, tumors and TDLN were harvested and manually dissociated. Blood was collected by submandibular vein collection, treated with RBC lysis buffer (BioLegend), and washed with phosphate-buffered saline prior to staining. Cells were treated with CD16/32 antibody (BioLegend) to prevent non-specific binding. Live cell staining was performed using Ghost Red Dy 780 (Tonbo Biosciences) according to manufacturer’s instruction. After live-dead staining, a single cell suspension was labeled with the surface antibodies at 4°C for 30 min and washed using filtered flow buffer (2% FBS + 2 mM EDTA in PBS). For antigen-specific CD8+ T cell staining, surface staining was performed with a MOC2 tumor specific peptide KSPWFTTL-H-2K^b^ tetramer (KSPWFTTL-PE, MBL) (CITE). For intracellular staining, cells were fixed and stained for internal markers with permeabilization solution according to manufacturer’s instructions (Cytek FOXP3 / Transcription Factor Staining Buffer Kit). Flow cytometry was performed using an Attune NxT Flow Cytometer (ThermoFisher). Data was analyzed using FlowJo Software. A complete list of antibody targets, clones, and fluorophores is provided in **Table S1**. Number of animals per group is indicated in figure legends.

### Spatial transcriptomics

Visium HD spatial transcriptomic sequencing data were processed using Space Ranger (v4.1; 10X Genomics) and aligned to the mouse reference genome (GRCm39). Subsequent analyses were performed in R using Seurat (v5.5.1). Following quality-control filtering, 8-μm spatial bins from all MOC2 tumors were used to construct a balanced reference dataset containing 50,000 bins per specimen (200,000 total bins). Principal Component Analysis (PCA), shared-nearest-neighbor, and unsupervised clustering were performed to identify transcriptionally distinct tissue states, which were visualized using Uniform Manifold Approximation and Projection (UMAP). Cluster identities were assigned using differentially expressed and canonical lineage markers and subsequently projected onto the full-resolution spatial datasets for visualization and quantification of tissue-state composition.

Regional biological responses associated with radiation exposure were evaluated using transcriptional programs representing VCAM1-associated endothelial adhesion, type I interferon (IFN1) signaling, ad MHCI antigen presentation. Response scores were standardized relative to the ICI treated tumor, with individual 8-μm binds exceeding 1 SD above the ICI reference mean considered positive for the corresponding program. To generate mutually exclusive composite response states, bins were hierarchically classified as Baseline, Low (VCAM1), Moderate (IFN1), or High (MHCI), with MHCI overriding IFN1 and IFN1 overriding VCAM1. Classified bins were subsequently aggregated into fixed 64-μm spatial neighborhoods to visualize regional response heterogeneity. Standardized VCAM1, IFN1, and MHCI axes were additionally analyzed independently across pairwise Low, Intermediate, and High response-state combinations.

Immune-associated spatial bins were isolated from the whole-tissue dataset and re-clustered to resolve major immune transcriptional states. Refined CD8-priming DC-like candidate bins were identified using combined DC-lineage, MHCII/APC, and cDC1/cross-presentation transcriptional features, with a migration-associated subset further characterized using *Ccr7*, *Fscn1*, *Relb*, and *Lamp3*. Spatial localization enrichment of migration-positive DC-like candidate was calculated as the percentage of candidate bins occupying a given response state minus the percentage of available tissue represented by that state. Localization was evaluated across both composite response categories and independent pairwise response axes, with statistical significance assessed using spatial-block permutation testing (nominal 256-μm blocks) and Benjamini-Hochberg false-discovery-rate correction. Candidate T-like candidate bins were conservatively identified through co-detection of CD8-lineage and T-cell-associated transcripts, with *Gzmb* and *Ifng* expression used to characterize cytotoxic and cytokine-associated effector states, respectively. Population abundance was normalized to the total number of QC-retained 8-μm tissue bins within each specimen.

### Statistical Analysis

Prism 10 (GraphPad Software) was used to create figures and calculate t-tests and ANOVA. Mann-Whitney t-test was used for two-group comparisons. Two-way ANOVA with Tukey’s honestly significant difference (HSD) test to adjust for multiple comparisons was used to assess statistical significance of observed mean differences for flow cytometry. One sample t-test was used to assess the statistical significance of the cytokine production fold change. The time-weighted average tumor volume of each mouse was calculated from all available data using the trapezoidal method. For the MOC2 experiment, ANOVA was used to test for any difference in the time-weighted average tumor volumes between the 4 treatment groups. Since the overall test was significant, pairwise t-tests were also used to compare specific groups. Since the other experiment only had 2 groups, a single t-test was used to compare dual ICI alone and the combination of dual ICI with ^177^Lu-NM600. Overall survival was plotted using the Kaplan-Meier method and compared using log-rank tests. P-values are presented without adjustment for multiple comparisons. Analysis was carried out in SAS 9.4.1.

## Author Contribution

MET, YW, WJJ, PAC: animal work and analysis. MET, WJJ, PAC, YW: flow cytometry staining and analysis. MBI, RTH: radiochemistry. MET, PC: RPT toxicity studies. OK, CDH, BPB: SPECT analysis and ^177^Lu-NM600 dosimetry. AOA, OK, BPB: iQID acquisition and analysis. MET and MH: biostatistical analysis. MET, ZD, IO: bioinformatics analysis. MET and ZSM: experimental design. JW, ZSM: funding, planning, and supervision of the project and data interpretation. MET, TB, ZSM wrote the manuscript. All authors read and approved the final manuscript.

## Acknowledgements

The authors would like to thank the University of Wisconsin Carbone Cancer Center (UWCCC) for supporting this project. The authors would like to thank Dr. Willliam Bachovchin from Tufts university for providing the FAP engineered cell lines. The authors would like to thank Point Biopharma Eli Lily and Archeus for providing key radiopharmaceutical agents.

