## Supplemental Data for "Optimized dosing of spatially heterogeneous radiopharmaceutical therapy reprograms poorly immunogenic tumors to enhance response to immune checkpoint blockade"

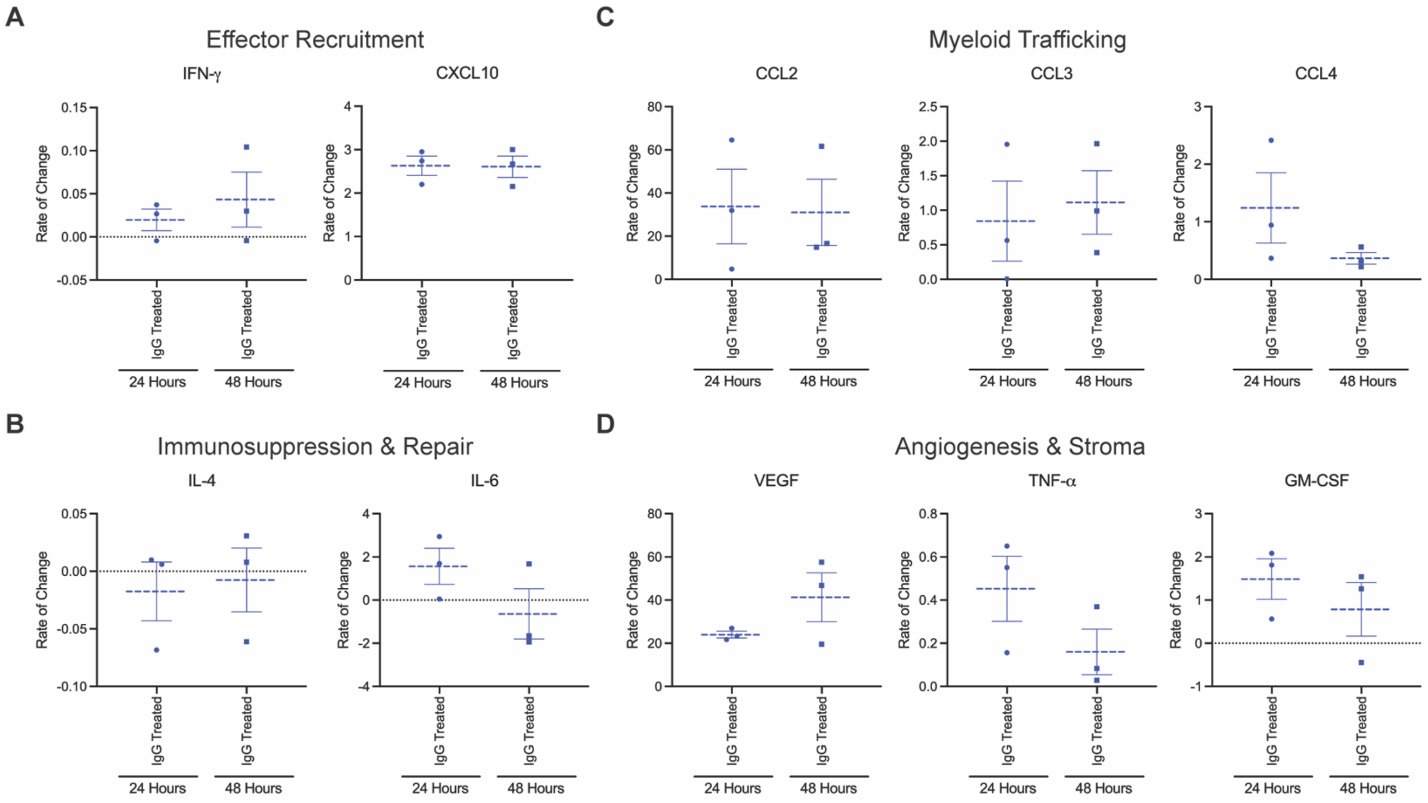


**Supplementary Figure 1. Cytokine and chemokine profile of IgG treated MOC2 cores.** A-D) Untreated MOC2 tumor-bearing mice. Tumors were harvested at day 12 and core needle biopsies were taken *ex vivo* and cultured using the Elephas Live Edge workflow. A) Analysis of IFN-γ and CXCL10 rate of change. B) Analysis of IL-4 and IL-6 rate of change. C) Analysis of CCL2, CCL3, and CCL4 rate of change. D) Analysis of VEGF, GM-CSF, and TNF-α rate of change.

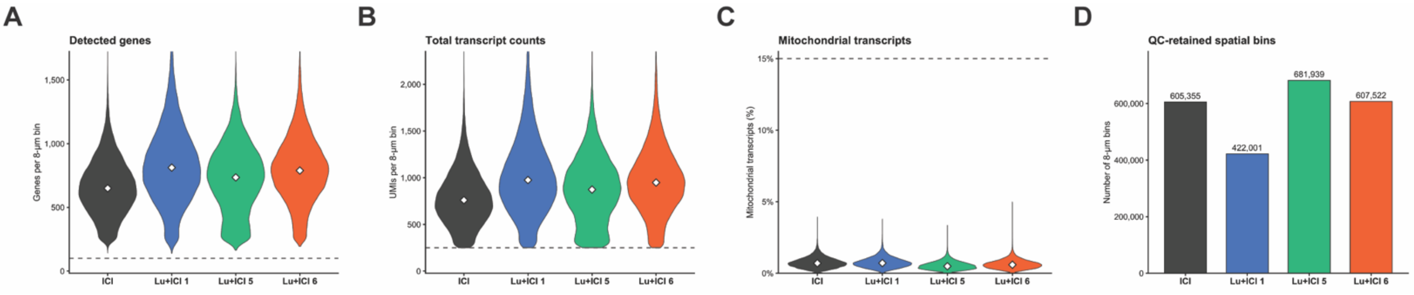


**Supplementary Figure 2. Quality control metrics of Visium HD spatial transcriptomic data.** A-D) Visium HD spatial transcriptomic analysis of MOC2 tumors treated with dual ICI (n=1) or $^{177}$Lu-NM600 + dual ICI (n=3). A) Distribution of detected genes per QC-retained 8-μm spatial bin. B) Distribution of total transcript counts per QC-retained 8-μm spatial bin. C) Percentage of mitochondrial transcripts across spatial bins, with the dashed line indicating the 15% QC threshold. D) Total number of QC-retained 8-μm spatial bins for each specimen.

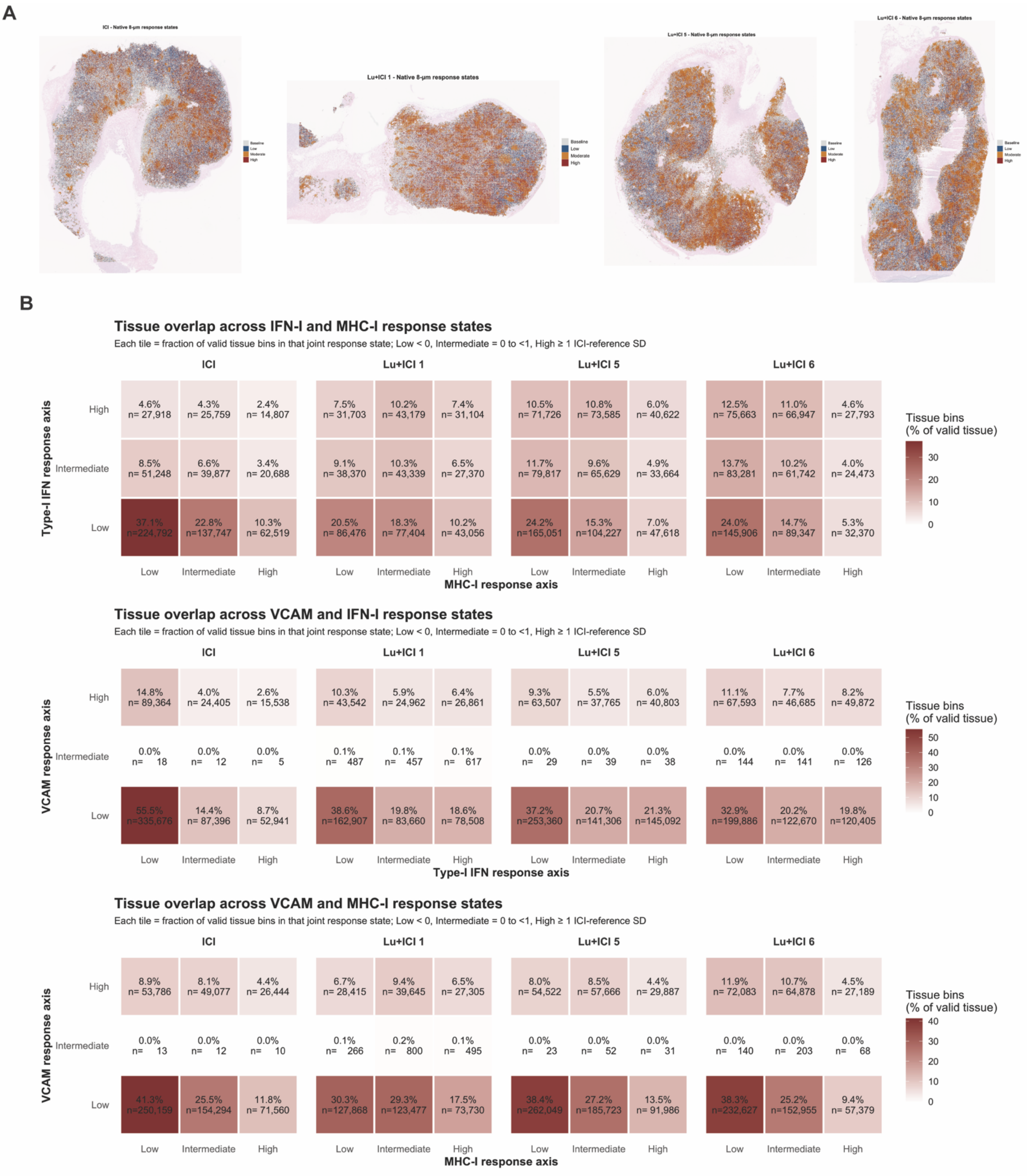
A
ICI - Native 8-µm response states
Lu+ICI 1 - Native 8-µm response states
Lu+ICI 5 - Native 8-µm response states
Lu+ICI 6 - Native 8-µm response states
Baseline
Low
Moderate
High
B
Tissue overlap across IFN-I and MHC-I response states
Each tile = fraction of valid tissue bins in that joint response state; Low < 0, Intermediate = 0 to <1, High ≥ 1 ICI-reference SD
ICI
Lu+ICI 1
Lu+ICI 5
Lu+ICI 6
Type-I IFN response axis
High
Intermediate
Low
4.6% n= 27,918
4.3% n= 25,759
2.4% n= 14,807
7.5% n= 31,703
10.2% n= 43,179
7.4% n= 31,104
10.5% n= 71,726
10.8% n= 73,585
6.0% n= 40,622
12.5% n= 75,663
11.0% n= 66,947
4.6% n= 27,793
8.5% n= 51,248
6.6% n= 39,877
3.4% n= 20,688
9.1% n= 38,370
10.3% n= 43,339
6.5% n= 27,370
11.7% n= 79,817
9.6% n= 65,629
4.9% n= 33,664
13.7% n= 83,281
10.2% n= 61,742
4.0% n= 24,473
37.1% n=224,792
22.8% n=137,747
10.3% n= 62,519
20.5% n= 86,476
18.3% n= 77,404
10.2% n= 43,056
24.2% n=165,051
15.3% n=104,227
7.0% n= 47,618
24.0% n=145,906
14.7% n= 89,347
5.3% n= 32,370
Low
Intermediate
High
MHC-I response axis
Tissue bins (% of valid tissue)
30
20
10
0
Tissue overlap across VCAM and IFN-I response states
Each tile = fraction of valid tissue bins in that joint response state; Low < 0, Intermediate = 0 to <1, High ≥ 1 ICI-reference SD
ICI
Lu+ICI 1
Lu+ICI 5
Lu+ICI 6
VCAM response axis
High
Intermediate
Low
14.8% n= 89,364
4.0% n= 24,405
2.6% n= 15,538
10.3% n= 43,542
5.9% n= 24,962
6.4% n= 26,861
9.3% n= 63,507
5.5% n= 37,765
6.0% n= 40,803
11.1% n= 67,593
7.7% n= 46,685
8.2% n= 49,872
0.0% n= 18
0.0% n= 12
0.0% n= 5
0.1% n= 487
0.1% n= 457
0.1% n= 617
0.0% n= 29
0.0% n= 39
0.0% n= 38
0.0% n= 144
0.0% n= 141
0.0% n= 126
55.5% n=335,676
14.4% n= 87,396
8.7% n= 52,941
38.6% n=162,907
19.8% n= 83,660
18.6% n= 78,508
37.2% n=253,360
20.7% n=141,306
21.3% n=145,092
32.9% n=199,886
20.2% n=122,670
19.8% n=120,405
Low
Intermediate
High
Type-I IFN response axis
Tissue bins (% of valid tissue)
50
40
30
20
10
0
Tissue overlap across VCAM and MHC-I response states
Each tile = fraction of valid tissue bins in that joint response state; Low < 0, Intermediate = 0 to <1, High ≥ 1 ICI-reference SD
ICI
Lu+ICI 1
Lu+ICI 5
Lu+ICI 6
VCAM response axis
High
Intermediate
Low
8.9% n= 53,786
8.1% n= 49,077
4.4% n= 26,444
6.7% n= 28,415
9.4% n= 39,645
6.5% n= 27,305
8.0% n= 54,522
8.5% n= 57,666
4.4% n= 29,887
11.9% n= 72,083
10.7% n= 64,878
4.5% n= 27,189
0.0% n= 13
0.0% n= 12
0.0% n= 10
0.1% n= 266
0.2% n= 800
0.1% n= 495
0.0% n= 23
0.0% n= 52
0.0% n= 31
0.0% n= 140
0.0% n= 203
0.0% n= 68
41.3% n=250,159
25.5% n=154,294
11.8% n= 71,560
30.3% n=127,868
29.3% n=123,477
17.5% n= 73,730
38.4% n=262,049
27.2% n=185,723
13.5% n= 91,986
38.3% n=232,627
25.2% n=152,955
9.4% n= 57,379
Low
Intermediate
High
MHC-I response axis
Tissue bins (% of valid tissue)
40
30
20
10
0

**Supplementary Figure 3. Native spatial distribution and overlap of radiation-associated transcriptional response states.** A-B) Visium HD spatial transcriptomic analysis of MOC2 tumors treated with dual ICI (n=1) or $^{177}$Lu-NM600 + dual ICI (n=3). A) Spatial distribution of native 8-μm tissue bins classified into Baseline, Low (VCAM1), Moderate (IFN1), High (MHCI) response states. B) Independent pairwise analysis of IFN1 x MHCI (top), VCAM1 x IFN1 (middle), and VCAM1 x MHCI (bottom) response programs across specimens. Each response axis was categorized as Low (below 0), Intermediate (0 to less than 1), or High (at least 1 ICI-reference SD). Heatmap values represent the percentage of valid tissue occupying each joint response state, with corresponding bin counts indicated.

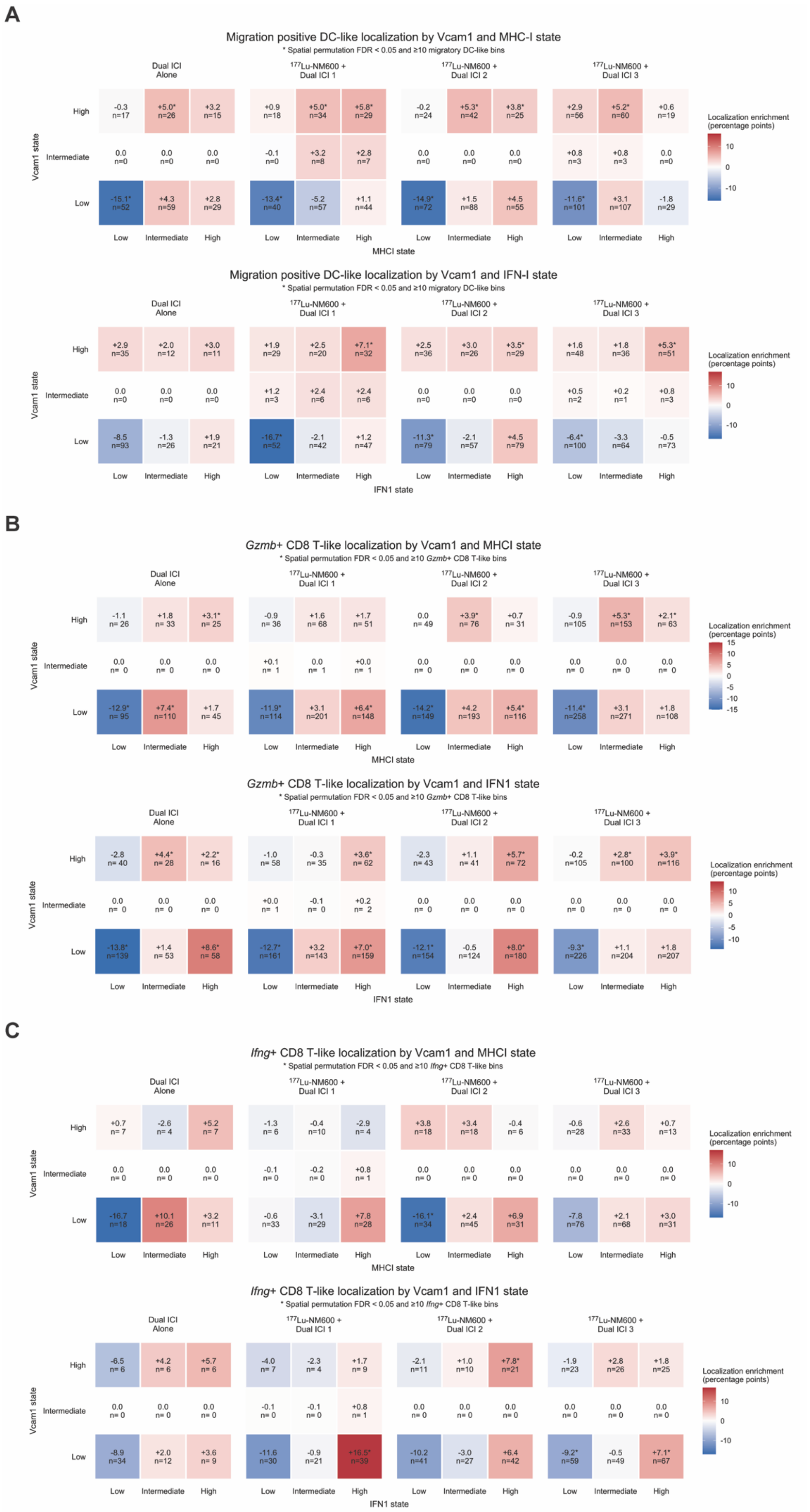
A
Migration positive DC-like localization by Vcam1 and MHC-I state
* Spatial permutation FDR < 0.05 and ≥10 migratory DC-like bins
Dual ICI Alone
177Lu-NM600 + Dual ICI 1
177Lu-NM600 + Dual ICI 2
177Lu-NM600 + Dual ICI 3
Vcam1 state
High
Intermediate
Low
MHCI state
Localization enrichment (percentage points)
Migration positive DC-like localization by Vcam1 and IFN-I state
* Spatial permutation FDR < 0.05 and ≥10 migratory DC-like bins
IFN1 state
B
Gzmb+ CD8 T-like localization by Vcam1 and MHCI state
* Spatial permutation FDR < 0.05 and ≥10 Gzmb+ CD8 T-like bins
Gzmb+ CD8 T-like localization by Vcam1 and IFN1 state
C
Ifng+ CD8 T-like localization by Vcam1 and MHCI state
* Spatial permutation FDR < 0.05 and ≥10 Ifng+ CD8 T-like bins
Ifng+ CD8 T-like localization by Vcam1 and IFN1 state

**Supplementary Figure 4. Spatial localization of migratory DC-like candidates and CD8 T-like effector states across MOC2 tumors.** A-C) Visium HD spatial transcriptomic analysis of MOC2 tumors treated with dual ICI (n=1) or $^{177}$Lu-NM600 + dual ICI (n=3). A) Localization enrichment of migration-positive DC-like candidate bins across joint VCAM1 x MHCI (top) and VCAM1 x IFN1 (bottom) response states. B) Localization enrichment of *Gzmb*+ T-like candidate bins across joint VCAM1 x MHCI (top) and VCAM1 x IFN1 (bottom) response states. C) Localization enrichment of *Ifng*+ T-like candidate bins across joint VCAM1 x MHCI (top) and VCAM1 x IFN1 (bottom) response states. Values represent enrichment relative to available tissue (percentage points), with candidate-bin counts indicated; *BH-adjusted spatial premutation FDR < 0.05 with ≥10 candidate bins (A-C).

**Table S1.** List of flow cytometry antibody targets, clones and fluorophores

| Name | Clone | Fluorophore | Catalog number |
|---|---|---|---|
| CD3 | 17A2 | FITC | BioLegend 100204 |
| CD8a | S18018E | PerCP-eFluor710 | BioLegend 162310 |
| FOXP3 | MF-14 | PE | BioLegend 126404 |
| NK1.1 | S17016D | PE-Dazzle594 | BioLegend 156518 |
| CD45 | S18009F | PE-Cy7 | BioLegend 157206 |
| MHC-I (H-2K$^b$) | AF6-88.5 | BV421 | BioLegend 116525 |
| CD4 | GK1.5 | BV510 | BioLegend 100449 |
| CD11c | N418 | BV605 | BioLegend 117334 |
| B220 | RA3-6B2 | BV711 | BioLegend 103255 |
| CD25 | 3C7 | APC | BioLegend 101910 |
| CD11b | M1/70 | FITC | BioLegend 101206 |
| CD11c | N418 | PerCP-Cy5.5 | BioLegend 117328 |
| CD172a | P84 | PE | BioLegend 144012 |
| XCR1 | ZET | PE-Dazzle594 | BioLegend 148234 |
| CD45 | S18009F | PE-Cy7 | BioLegend 157206 |
| CD86 | PO3 | BV421 | BioLegend 105123 |
| MHCII (I-A/I-E) | M5/114.15.2 | BV510 | BioLegend 107636 |
| CD103 | 2E7 | BV605 | BioLegend 121433 |
| CD80 | 16-10A1 | BV711 | BioLegend 104743 |
| Siglec H | 551 | Alexa647 | BioLegend 129608 |
| B220 | RA3-6B2 | Alexa700 | BioLegend 103232 |
